# FAM210A Regulates Mitochondrial Translation and Maintains Cardiac Mitochondrial Homeostasis

**DOI:** 10.1101/2023.05.20.541585

**Authors:** Jiangbin Wu, Kadiam C Venkata Subbaiah, Omar Hedaya, Si Chen, Joshua Munger, Wai Hong Wilson Tang, Chen Yan, Peng Yao

## Abstract

**Aims:** Mitochondria play a vital role in cellular metabolism and energetics and support normal cardiac function. Disrupted mitochondrial function and homeostasis cause a variety of heart diseases. Fam210a (family with sequence similarity 210 member A), a novel mitochondrial gene, is identified as a hub gene in mouse cardiac remodeling by multi-omics studies. Human FAM210A mutations are associated with sarcopenia. However, the physiological role and molecular function of FAM210A remain elusive in the heart. We aim to determine the biological role and molecular mechanism of FAM210A in regulating mitochondrial function and cardiac health *in vivo*.

**Methods and Results:** Tamoxifen-induced *αMHC*^MCM^-driven conditional knockout of *Fam210a* in the mouse cardiomyocytes induced progressive dilated cardiomyopathy and heart failure, ultimately causing mortality. Fam210a deficient cardiomyocytes exhibit severe mitochondrial morphological disruption and functional decline accompanied by myofilament disarray at the late stage of cardiomyopathy. Furthermore, we observed increased mitochondrial reactive oxygen species production, disturbed mitochondrial membrane potential, and reduced respiratory activity in cardiomyocytes at the early stage before contractile dysfunction and heart failure. Multi-omics analyses indicate that FAM210A deficiency persistently activates integrated stress response (ISR), resulting in transcriptomic, translatomic, proteomic, and metabolomic reprogramming, ultimately leading to pathogenic progression of heart failure. Mechanistically, mitochondrial polysome profiling analysis shows that FAM210A loss of function compromises mitochondrial mRNA translation and leads to reduced mitochondrial encoded proteins, followed by disrupted proteostasis. We observed decreased FAM210A protein expression in human ischemic heart failure and mouse myocardial infarction tissue samples. To further corroborate FAM210A function in the heart, AAV9-mediated overexpression of FAM210A promotes mitochondrial-encoded protein expression, improves cardiac mitochondrial function, and partially rescues murine hearts from cardiac remodeling and damage in ischemia-induced heart failure.

**Conclusion:** These results suggest that FAM210A is a mitochondrial translation regulator to maintain mitochondrial homeostasis and normal cardiomyocyte contractile function. This study also offers a new therapeutic target for treating ischemic heart disease.

**Translational Perspective:** Mitochondrial homeostasis is critical for maintaining healthy cardiac function. Disruption of mitochondrial function causes severe cardiomyopathy and heart failure. In the present study, we show that FAM210A is a mitochondrial translation regulator required for maintaining cardiac mitochondrial homeostasis *in vivo*. Cardiomyocyte-specific FAM210A deficiency leads to mitochondrial dysfunction and spontaneous cardiomyopathy. Moreover, our results indicate that FAM210A is downregulated in human and mouse ischemic heart failure samples and overexpression of FAM210A protects hearts from myocardial infarction induced heart failure, suggesting that FAM210A mediated mitochondrial translation regulatory pathway can be a potential therapeutic target for ischemic heart disease.

## Introduction

The mitochondrion is an essential cellular organelle to maintain the energy supply for any biochemical reactions in eukaryotic cells ^1^. The electron transport chain (ETC) complex, located in the mitochondrial inner membrane, generates an electrochemical-proton gradient that drives ATP synthesis. In addition to energetics, mitochondria also play critical roles in calcium signaling, metabolism, and apoptosis. The heart is one of the most mitochondria-rich organs in mammals, as mitochondria account for over 30% of cardiomyocyte (CM) cell mass ^2^. Mitochondria generate metabolites and ATP via tricarboxylic acid cycle (TCA) and oxidative phosphorylation, fueling all biological processes, including CM contraction. Loss of function of mitochondrial genes due to genetic mutations can cause mitochondrial cardiomyopathy (MC) ^3^ and other mitochondrial diseases in the skeletal muscle, brain, liver, and kidney, among other organs ^4^. Moreover, reduced expression of critical mitochondrial proteins and subsequent mitochondrial dysfunction are major drivers of pathogenesis in ischemic heart disease such as myocardial infarction (MI) ^5, 6^. However, the etiology and effective therapeutic intervention of mitochondria dysfunction related heart disease are still lacking ^7, 8^.

Given these crucial roles of mitochondria in CMs, elucidating the molecular functions of critical regulators of mitochondrial homeostasis may provide novel therapeutic targets to combat MC and MI. FAM210A (family with sequence similarity 210 member A) is a mitochondria-localized protein essential for embryonic development and viability ^9^. Genome-wide association studies have shown multiple FAM210A mutations associated with sarcopenia and osteoporosis ^10, 11^. However, FAM210A is not expressed in the bone but the skeletal muscle and is most highly expressed in the heart ^11, 12^. Skeletal muscle-specific knockout (KO) of *Fam210a* reduces muscle strength and bone function *in vivo* ^11^, and reduced expression of FAM210A impairs myoblast differentiation *in vitro* ^13^. One unbiased transcriptome-wide RNA-seq and ribosome profiling analysis in mouse hearts identified FAM210A as a hub gene downregulated in cardiac pathological hypertrophy and remodeling compared to physiological hypertrophy ^14^. Recently, our study revealed that the miR-574-FAM210A axis plays a vital role in regulating mitochondrial proteomic homeostasis and cardiac remodeling ^12^. FAM210A interacts with a critical mitochondrial translation elongation factor EF-Tu in a protein complex involved in regulation of mitochondrial-encoded mRNA translation ^12^. However, the *in vivo* physiological function and downstream molecular effector of FAM210A in the heart remain unknown.

Here, we established a novel CM-specific Fam210a conditional KO mouse model to investigate the function of FAM210A in the CM and heart. We observed severe morphological disruption and functional decline in mitochondria, CMs, and the heart at ∼10 weeks post inducible *Fam210a* KO. Meanwhile, we only detected reduced mitochondrial membrane potential (Δλφλ_m_) and increased reactive oxygen species (ROS) accompanied by aberrant mitochondrial function in the absence of any dysfunction in CMs and the heart at the early stage (∼5 weeks post *Fam210a* KO). Multi-omics analyses in cKO hearts uncovered chronic activation of the integrated stress response (ISR) as a major component of the gene signature and disease etiology accompanied by transcriptomic, translatomic, proteomic, and metabolomic reprogramming. Mechanistically, mitochondrial polysome profiling analyses show that FAM210A modulates mitochondrial encoded mRNA translation and maintains mitochondrial homeostasis. FAM210A deficiency impairs mitochondrial proteomic homeostasis. Overexpression of FAM210A protects the hearts from cardiac injury and pathological remodeling in a mouse MI model. This study demonstrates that FAM210A maintains mitochondrial homeostasis in CMs by regulating the translation of mitochondria-encoded mRNAs and can be targeted for cardiac protection under ischemic stresses.

## Methods

A detailed description of materials and methods is available in *the* supplemental information.

### Mice

All mouse experiments were conducted under protocols approved by the University Committee on Animal Resources (UCAR) of the University of Rochester Medical Center (URMC). All animal procedures conform to the guidelines from Directive 2010/63/EU of the European Parliament on the protection of animals used for scientific purposes or the NIH Guide for the Care and Use of Laboratory Animals. The mice were housed in a 12:12 hours light: dark cycle in a temperature-controlled room in the animal housing room of URMC, with free access to water and food. The age and gender were indicated below in each section of experiments using mice. The *Fam210a* floxed mice (tm1c, *Fam210a*^flox/flox^) were generated by the Canadian Mouse Mutant Repository (CMMR, http://www.cmmr.ca/) and purchased through the International Mouse Phenotyping Consortium (https://www.mousephenotype.org/). The tamoxifen-inducible transgenic mouse line *αMHC*^MerCreMer^ (*αMHC*^MCM/+^) was a gift from Dr. Eric Small’s lab at URMC. The tamoxifen-induced cardiomyocyte-specific knockout of *Fam210a* was achieved by crossing *Fam210a*^flox/flox^ *mice with αMHC*^MCM/+^ mice. Age and gender-matched *aMHC*^MCM/+^ mice were used to control *Fam210a* knockout mice. All the mice are on C57BL/6J background.

Anesthetic and analgesic agents used in the study are listed below:

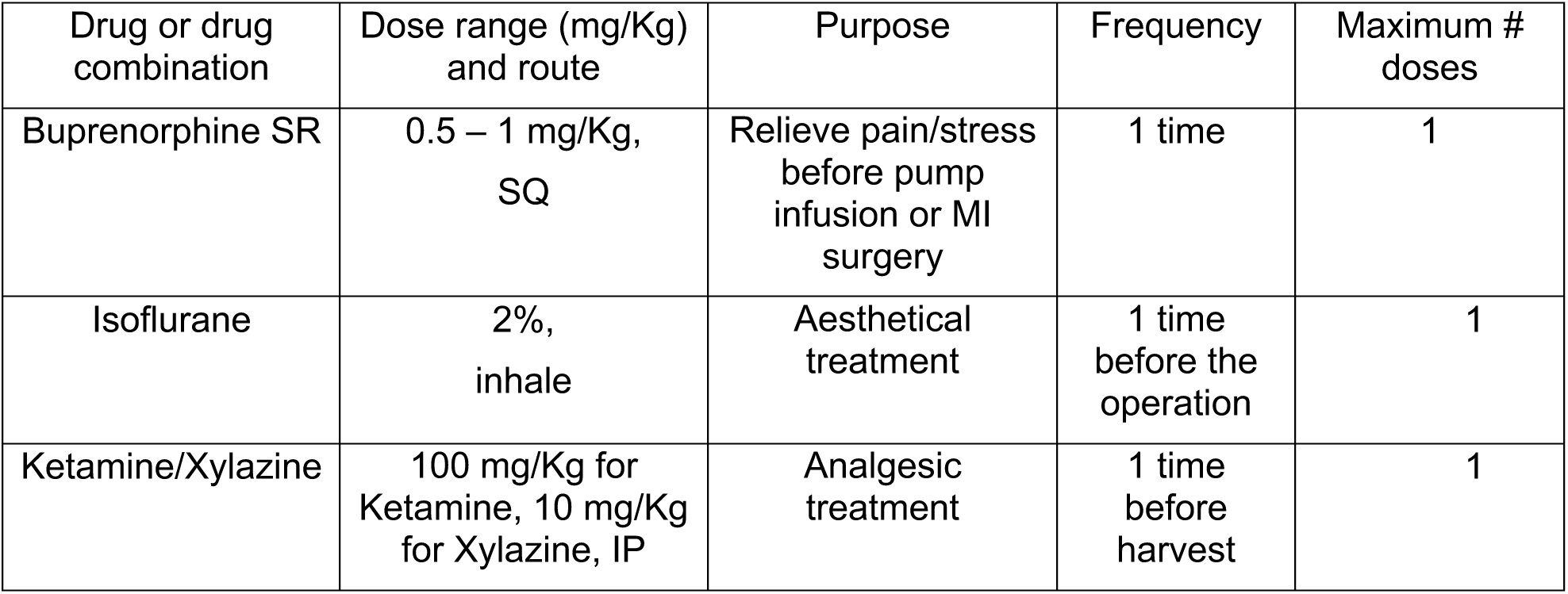

After Buprenorphine SR treatment, lab fellows inspected mice once a day for 3 days and registered them in the recording card. Carbon dioxide euthanasia is used for terminating mice. All anesthetic and analgesic agents used in the study were approved by the UCAR of the URMC.

### Statistical Analysis

All quantitative data were presented as mean ± SEM and analyzed using Prism 8.3.0 software (GraphPad). Kolmogorov-Smirnov test was used to calculate the normal distribution of the data. For normally distributed data, an unpaired two-tailed Student t-test was performed to compare two groups and one-way or two-way ANOVA with Tukey’s multiple comparisons test for the comparisons between more than three groups. For data that is not normally distributed, a non-parametric Mann-Whitney test was performed for the comparisons between two groups and a Kruskal-Wallis test with Dunn’s multiple comparisons test for the comparisons between more than three groups. The Log-rank test was performed to compare survival rates, and the χ^2^ test was used to compare cell number counting data. Two-sided *P* values <0.05 were considered to indicate statistical significance. Specific statistical methods were described in the figure legends.

## Results

### Cardiomyocyte-specific Fam210a knockout causes spontaneous dilated cardiomyopathy

To investigate the role of FAM210A *in vivo*, we first determined the expression of FAM210A in two major cell types in the heart, CM and cardiac fibroblast (CF). FAM210A is highly expressed in CMs compared with CFs (Figure S1A). Moreover, single-cell RNA-seq data reveals that the highest expression is found in CMs than in any other human cell type (Figure S1B) ^15^. To determine the role of FAM210A in the adult heart, we generated an inducible CM-specific cKO mouse model of *Fam210a* by cross-breeding *Fam210a*^flox/flox^ mice with tamoxifen (TMX)-inducible *αMHC*^MCM/+^ mice (Figure 1A, S1C, D). After TMX administration, FAM210A mRNA and protein expression was decreased by >90% in the cKO heart (Figure S1E, F). The cKO mice declined in bodyweight 9 weeks post-TMX induced KO (Figure 1B). Both male and female cKO mice showed complete penetrance of lethality after TMX injection for ∼70-80 days (Figure 1C), with a life span of 77.20 ± 1.07 days in females and 74.89 ± 1.58 days in males. Echocardiographic examinations suggest that *Fam210a* cKO mice experienced progressive dilated cardiomyopathy with decreased fractional shortening, ejection fraction (Figure 1D), and induced chamber dilation (Figure S1G-J, Table S1) compared to *αMHC*^MCM/+^ (Ctrl) and *Fam210a* fl/fl mice. As we did not observe any significant difference in cardiac functions between *αMHC*^MCM/+^ and *Fam210a* fl/fl mice in the echocardiographic outcome within 11 weeks post TMX injections (Table S1), we used *αMHC*^MCM/+^ mice as the control group in the following phenotyping and multi-omics studies. The cardiac function remained normal until 7 weeks post KO and started to drop severely after ∼9 weeks. Therefore, we defined the early (1-7 weeks) and late stages (8-11 weeks) of this cardiomyopathy disease model using 7 weeks post KO as the cut-off time point of transition from normal to abnormal cardiac function. We first characterized the disease phenotypes at the late stage at ∼10 weeks post KO. The increased ratio of heart weight to tibia length (by 19.56% in males, by 18.93% in females) and myocyte cross-section area by wheat germ agglutinin (WGA) staining confirmed spontaneous dilated cardiomyopathy with hypertrophied CMs in cKO mice (Figure 1E, F). Consistent with the marked CM hypertrophy, hypertrophy marker gene expression (Myh7, Nppa, and Nppb) was drastically induced, while Myh6 expression was reduced in cKO hearts compared with control hearts (Figure 1G). Furthermore, a slight increase in cardiac fibrosis was indicated by Picrosirius red staining and confirmed by a modest rise in fibrosis marker gene expression (Col1a1 and Col3a1) (Figure 1H, I). Altogether, these results suggest that CM-specific *Fam210a* KO in adult mice leads to progressive HF with enlarged CMs and dilated left ventricle chamber, and ultimately causes mortality at ∼75 days after *Fam210a* KO.

**Figure 1.**
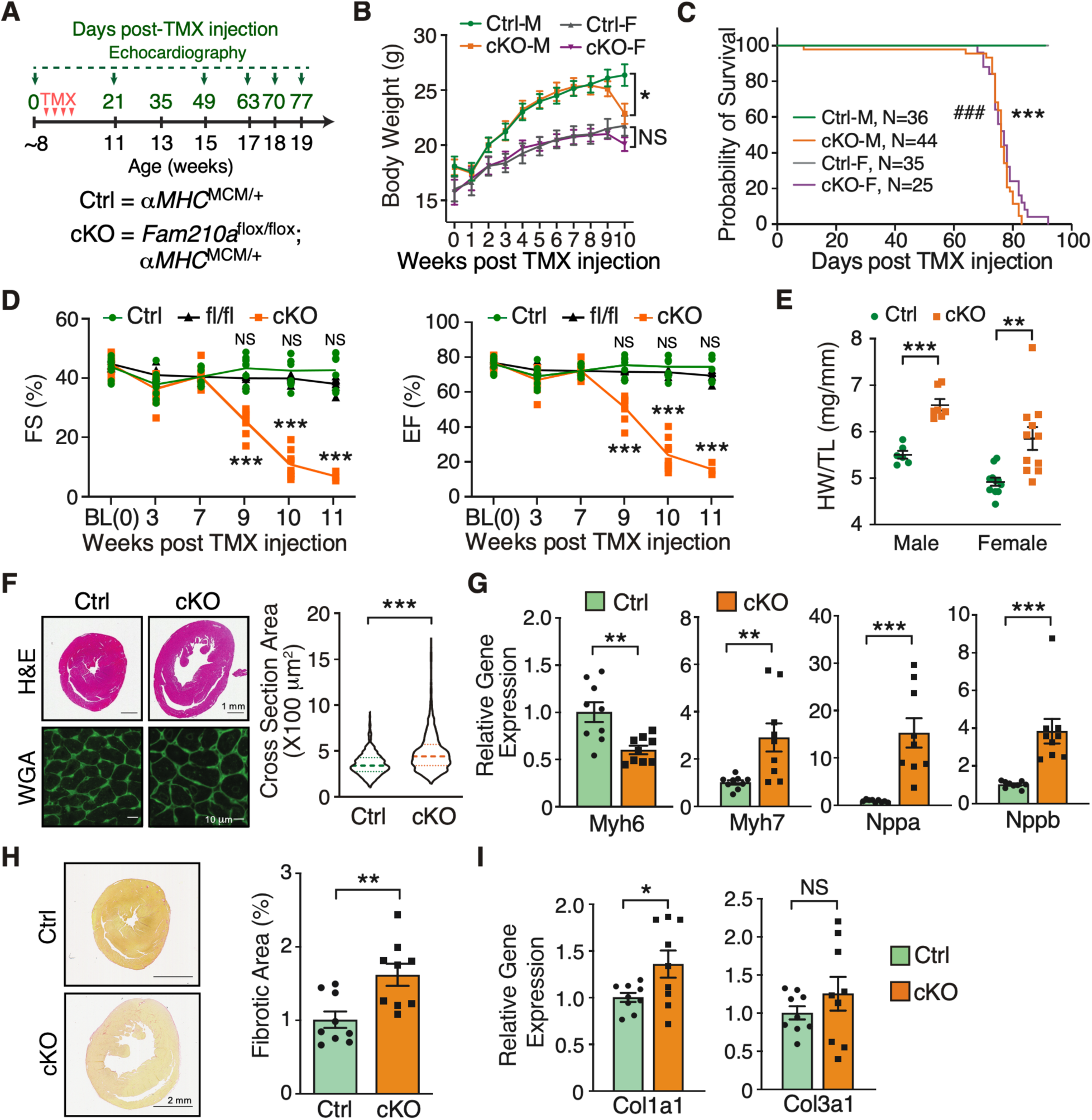
FAM210A deficiency in cardiomyocytes leads to dilated cardiomyopathy and heart failure. (**A**) Experimental timeline for phenotypic examination of CM-specific *Fam210a* cKO mice. (**B**) The body weight was measured weekly post-TMX-induced *Fam210a* KO in male and female mice. N = 11 for Ctrl and cKO male (M). N = 7 for Ctrl female (F) and N = 10 for cKO female. (**C**) The survival rate in male and female mice post KO. ### *P* < 0.001 for M and *** *P* < 0.001 for F by Log-rank test. (**D**) Fractional shortening (FS) and ejection fraction (EF) were measured by echocardiography in cKO and control mice. N = 5M + 4F for Ctrl and cKO. N = 5M + 2F for *Fam210a* fl/fl control mice. NS: not significant for fl/fl versus Ctrl by Two-way ANOVA with Sidak multiple comparisons. (**E**) Heart weight/Tibia length (HW/TL) ratio in male and female cKO mice. N = 6 and 7 for male Ctrl and cKO. N = 11 for female Ctrl and cKO. (**F, H**) Wheat germ agglutinin (WGA) staining for the cross-sectional area of CMs (F; N = 3 hearts with >1000 CMs quantified per heart) and picrosirius red staining of collagen deposition (H; N = 5M + 4F for Ctrl and cKO) in the hearts of control and cKO mice at ∼65 days post-KO. (**G, I**) Cardiac hypertrophy (G) and fibrosis (I) marker gene expression at ∼65 days post-KO (N = 5M + 4F). * *P* < 0.05; ** *P* < 0.01; *** *P* < 0.001 by two-way ANOVA with Sidak multiple comparisons test (B, D), student t-test (E, G, H, I), and Mann-Whitney test (F and Nppb in G).

### FAM210A deletion leads to myofilament disarray and contractile dysfunction of cardiomyocytes

To dissect the pathological effects of FAM210A deletion on CM cells, we first performed transmission electron microscopy to visualize the morphological changes in diseased CMs in the whole heart tissue sections of *Fam210a* cKO mice at the late stage of cardiomyopathy (∼10 weeks post KO). The sarcomeric architecture was remarkably disrupted, as indicated by the destruction of the organized Z disk, I band, M line (Figure 2A and S2A), and the loss of intercalated disc integrity between CMs in the cKO hearts (Figure S2B). The observation of myofilament disarray was confirmed in isolated primary CMs from cKO and control mice by immunostaining of Z disk and thin filament marker protein α-actinin 2 and cardiac troponin T, respectively (Figure 2B, C, S2C). In addition, quantitative mass spectrometry analysis was performed to evaluate the proteomic changes in the whole heart tissues of *Fam210a* cKO and control mice. The sarcomeric protein levels of Troponin C1, Troponin T2, Troponin I3, Myosin heavy chain 6 and 3, Myosin light chain 2, and Tropomyosin 1 (Log2FC = -0.66) were decreased while Myosin heavy chain 7, Myosin light chain 4 and 7, α-actinin 1 and 4, and Tropomyosin 2/3/4 (Log2FC = 0.95, 0.88, and 0.53) were increased (Figure S2D, Table S3), suggesting a proteomic homeostatic imbalance of myofilament proteins in cKO hearts. Immunoblot experiments verified the same trend of changes of α-actinin (increased by 2.66 folds) and cardiac Troponin T (decreased by 40.9%) in the whole heart lysate of cKO hearts (Figure S2E). To examine whether dysregulation of sarcomeric protein expression affects CM contractility, we measured the dynamic change in sarcomere length during the contraction cycle of CMs isolated from control and cKO mice at the late stage using the myocyte contractility recording system. In agreement with dysregulated sarcomeric structure proteins, *Fam210a* KO CMs exhibited significantly compromised contractile function compared to control CMs in the presence or absence of isoproterenol (ISO) stimulation (Figure 2D-F, S2F H). The sarcomere shortening (Figure 2D, E) and maximal shortening velocity (Figure S2F) were reduced in KO CMs. As a result, time to 50% sarcomere shortening was increased in KO CMs (Figure S2G). Meanwhile, we noticed that maximal relengthening velocity was reduced (Figure S2H), and time to 50% relaxation was increased accordingly (Figure 2F). Taken together, these results suggest that genetic deletion of *FAM210A* in CMs leads to dysregulated expression of myofilament proteins and pronounced disruption of the sarcomeric structure, thereby driving contractile dysfunction in CMs.

**Figure 2.**
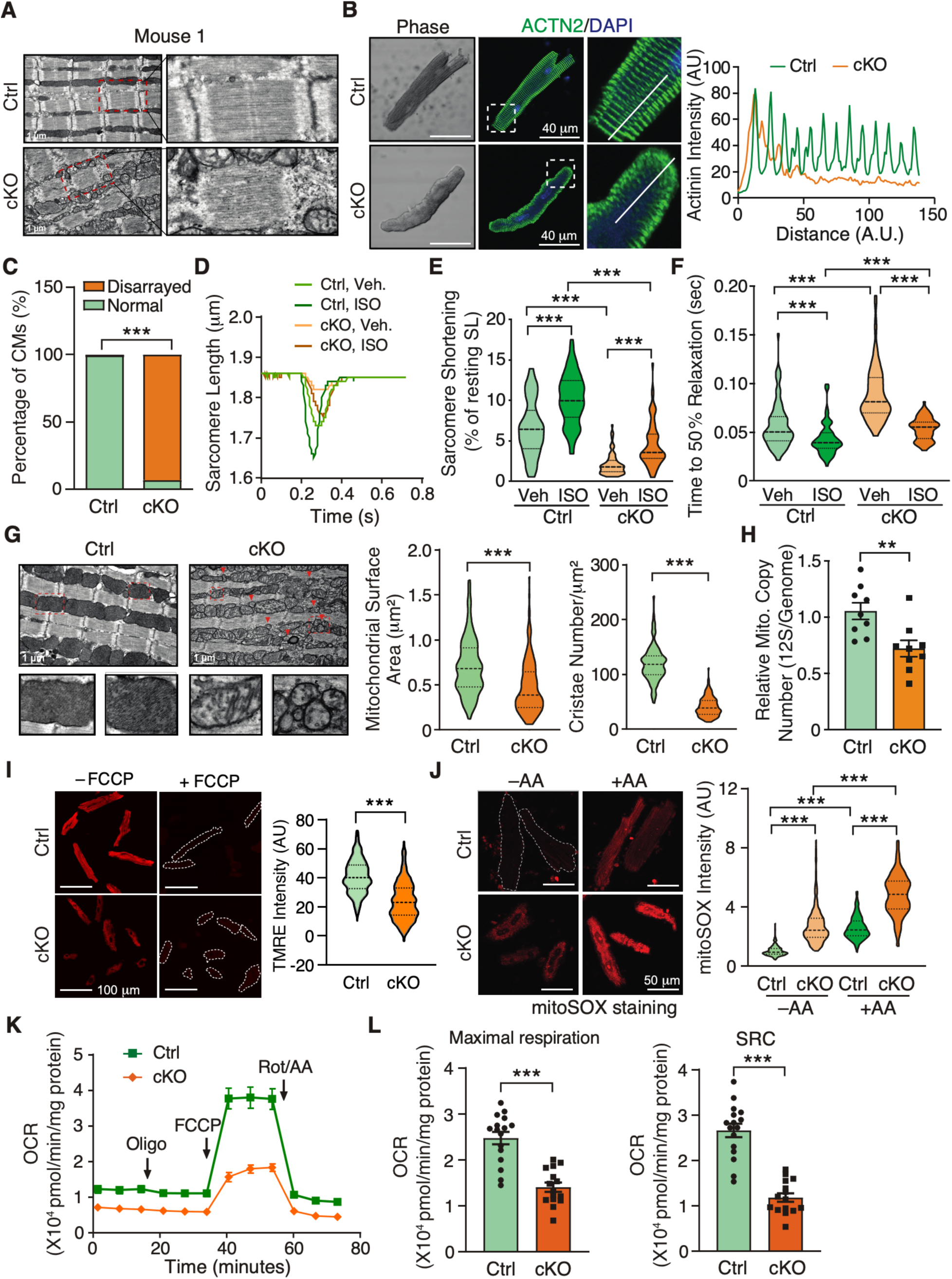
Cardiomyocyte-specific deletion of *Fam210a* causes myofilament disarray and mitochondrial dysfunction in cardiomyocytes. (**A**) A representative electron microscopy image shows a disturbed sarcomeric structure in CMs from whole heart tissue sections of a late-stage *Fam210a* cKO heart (∼65 days post-KO). (**B**) Immunofluorescence staining of ACTN2 indicates myofilament disarray in isolated CMs from cKO hearts at ∼65 days post-KO. Right panels: quantification of fluorescence intensity distribution (AU, arbitrary unit) along the white line in the IF image. (**C**) Quantification of disarrayed CM percentile from IF staining in isolated CMs from hearts of control or cKO mice. N > 2500 CMs were quantified from 4 hearts. (**D-F**) FAM210A deficiency reduces sarcomere length and attenuates the contractility of isolated CMs from cKO hearts at ∼65 days post-KO with or without ISO stimulation. N = 82/79/77/68 CMs were quantified for Ctrl-Veh, Ctrl-ISO, cKO-Veh, and cKO-ISO from 4 hearts. (**G**) Electron microscopy shows decreased cristae and mitochondrial size in *Fam210a* cKO CMs from whole heart tissue sections. Red triangles indicate fragmented mitochondria. The mitochondrial surface area and cristae number are quantified in >200 mitochondria from N = 3 hearts (cKO vs. Ctrl). (**H**) The mtDNA copy number in control and cKO hearts was measured by qPCR using primers targeting the mitochondrial 12S rDNA locus (N = 5M + 4F). qPCR of the nuclear genomic DNA was used as a normalizer. (**I**) Mitochondrial membrane potential (Δ\+l_m_) was determined by TMRE staining in isolated CMs from late-stage *Fam210a* cKO hearts. For the FCCP treatment, 10 μM FCCP was added into the medium for 5 min before imaging. >600 CMs from 4 hearts were quantified. The TMRE intensity was normalized by the intensity after FCCP treatment. (**J**) The ROS level in isolated CMs from late-stage cKO hearts. >200 CMs from 3 hearts were quantified. AA: antimycin A (2 mM for 30 min). (**K, L**) The mitochondrial respiratory activity in isolated CMs from control and cKO hearts at ∼65 days post-KO was measured by the Seahorse assay. N = 16 biological replicates for Ctrl and N = 15 for cKO from CM isolations in three hearts. SRC: spare respiratory capacity (OCR^Max^-OCR^Bas^). * *P* < 0.05, ** *P* < 0.01, *** *P* < 0.001 by χ^2^ test (C), Kruskal-Wallis test with Dunn’s multiple comparisons test (E, F, J), student t-test (H, L), and Mann-Whitney test (G, I).

### FAM210A maintains cardiac mitochondrial function in cardiomyocytes

Next, we ask the question of the effect of *Fam210a* KO on cardiac mitochondrial function. Electron microscopy imaging showed that mitochondrial morphology was dramatically altered in cKO hearts at the late stage of HF. Cristae were greatly reduced and sparsely present in the mitochondria of KO CMs compared to control CMs (Figure 2G, left panel). Also, increased number of small-sized mitochondria were observed in KO CMs but not in control CMs. Quantification results from electron microscopy images indicated that mitochondrial surface area and cristae number were reduced by 32.6% and 65.6%, respectively (Figure 2G, middle and right panels). In addition, the mitochondrial DNA (mtDNA) copy number was decreased by 31.60% and 36.67% as measured by PCR using specific primers targeting 12S rDNA or *Cox1* locus in the mitochondrial genome, respectively, in cKO hearts compared to control hearts at the late stage (Figure 2H and S3A). To determine the cause of reduced mtDNA copy number, we measured the expression of critical genes involved in mitochondrial biogenesis and mitophagy in the cKO hearts. Although *Pgc1α* and *Pgc1β* mRNA was decreased by 36.1% and 44.7%, respectively (Figure S3B), PGC1α protein expression was slightly increased, indicating that mitochondrial biogenesis may stay unchanged or modestly enhanced (Figure S3C). As a consequence of decreased mtDNA copy number in KO CMs, the protein expression of the mitochondrial localized transcription factor TFAM was reduced by half (Figure S3C). In other aspects, the expression of autophagy marker protein LC3B II was increased by ∼3-4 folds (Figure S3C) and the autophagic flux was enhanced in the presence of autophasome and lysosome fusion inhibitor, bafilomycin A1 (Baf A1), at the late stage (Figure S3D), suggesting that enhanced autophagic activity may contribute partially to the reduced mtDNA copy number at the late stage through mitochondria degradation. Furthermore, blue native gel assays showed that ETC assembly was compromised to some extent in *Fam210a* cKO hearts compared to control hearts at the late stage (Figure S3E). ETC assembly defect or dysfunction is often associated with compromised Δψ_m_ and increased mitochondrial ROS production ^16^. In isolated KO CMs, the Δψ_m_ was reduced by 37.3% (Figure 2I). ROS production was increased by 1.87 and 2.72 folds in the absence or presence of the Complex III inhibitor Antimycin A, respectively (Figure 2J). More importantly, seahorse measurement showed that the mitochondrial respiratory activity was significantly compromised in FAM210A deficient CMs (Figure 2K), as indicated by a ∼50% decrease in the maximal oxygen consumption rate and spare respiratory capacity (Figure 2L). Consistent with aberrant mitochondrial function, we observed a significantly increased (∼2.5%) CM cell death at the late stage of *Fam210a* cKO hearts (Figure S3F). These results suggest that FAM210A is required to maintain cardiac mitochondrial homeostasis, and its deficiency causes severe mitochondrial dysfunction and cardiomyopathy.

### Disrupted mitochondrial membrane potential and enhanced ROS production are early events in *Fam210a* cKO hearts

To tease out the disease-causing events from the consequent secondary effects of HF, we examined the functional changes in early-stage heart tissues of *Fam210a* cKO mice. We did not observe any functional abnormality of hearts and morphological change of CMs with a minimal expressional change of CM hypertrophy marker genes at the early stage (∼5 weeks post KO) (Figure 3A and S4A-C). Additionally, cardiac fibrosis was absent, and fibrosis marker gene expression was unaltered in cKO hearts (Figure S4D,E). Electron microscopy analysis indicated no significant disruption in the overall morphology of the CM sarcomeric network or mitochondrial structure (Figure 3B). Consistently, we did not identify any remarkable difference in contractile functions (Figure S4F-L) between KO and control CMs in the presence or absence of ISO stimulation. These results reveal no significant cardiac dysfunction in *Fam210a* cKO hearts at the early stage.

**Figure 3.**
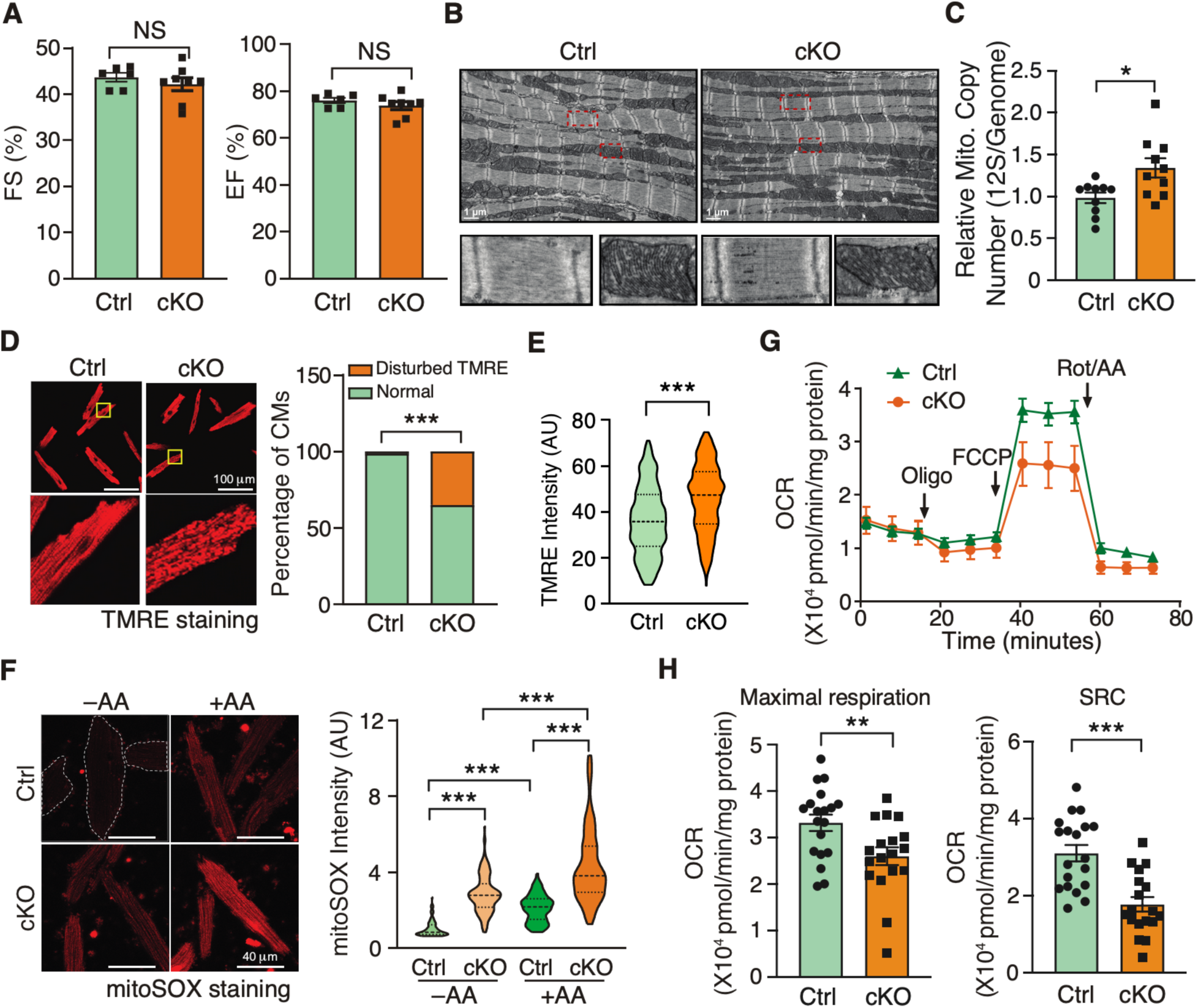
Disturbed mitochondrial membrane potential and increased ROS production are early events in FAM210A-deficient CMs. (**A-C**) Cardiac function (A, N = 6 for Ctrl and N = 8 for cKO) and mitochondrial morphology (B, N = 3) stay unchanged while mtDNA copy number is slightly increased as quantified by PCR using primers targeting 12S rDNA (C, N = 10 for Ctrl and cKO) at 5 weeks post-KO. (**D, E**) Mitochondrial membrane potential (Δψ_m_) was determined by TMRE staining in isolated CMs from early-stage *Fam210a* cKO hearts (D). Right panel: >1500 CMs were quantified from 4 hearts. For Δψ_m_ measurement with the FCCP treatment in (E), 10 μM FCCP was added into the medium for 5 min before imaging. >280 CMs were quantified from 4 hearts. The TMRE intensity was normalized by the intensity after FCCP treatment. Representative images are shown in Figure S5H. (**F**) Increased mitochondrial ROS (>80 CMs were quantified from 3 hearts) was observed in isolated CMs from early-stage cKO hearts. (**G, H**) Respiratory activity is attenuated in isolated CMs from early-stage KO hearts. N = 19 biological replicates for Ctrl and N = 18 for cKO in (G). Oligo: Oligomycin; FCCP: Carbonyl cyanide-p-trifluoromethoxyphenyl-hydrazone; Rot/AA: Rotenone/Antimycin A. * *P* < 0.05; ** *P* < 0.01; *** *P* < 0.001 by student t-test (A, C, H), χ^2^ test (D), Mann-Whitney test (E), and Kruskal-Wallis test with Dunn’s multiple comparisons test (F).

Given the localization of FAM210A in mitochondria, we assume that the primary disease-causing events are associated with mitochondria at the molecular level before any noticeable phenotypic changes occur in overall mitochondrial morphology, CMs, and the heart. To test this hypothesis, we first measured the mitochondrial surface area and cristae number and width, and noticed a slight increase in the cristae number and a modest decrease in the cristae width (Figure S5A). This altered cristae structure driven by loss of FAM210A can be partially explained by the previously reported interaction of FAM210A (C18ORF19) with ATAD3A ^17^ that controls cristae structure ^18^. However, we did not observe a significant change in ATAD3A protein expression in *Fam210a* cKO hearts at the early stage (Figure S5B). Additionally, given that mtDNA copy number was slightly increased at the early stage in cKO hearts compared to control hearts (Figure 3C and S5C), we measured the expression of mitochondria biogenesis factor genes and found that the expression of *Pgc1α* and *Pgc1β* mRNA was indeed reduced by 36.1% and 27.9%, respectively (Figure S5D). However, the protein expression of neither the nuclear nor mitochondrial localized mitochondria biogenesis factor, PGC1α and TFAM, was changed (Figure S5E). In contrast, autophagic activity was unchanged or slightly decreased as indicated by LC3B II protein expression with or without of Baf A1 treatment (Figure S5F), implying that autophagy-mediated mitochondria degradation pathway may have a minimal effect on the slightly increased mtDNA copy number at the early stage in *Fam210a* cKO hearts. Moreover, blue native gel assays showed that ETC assembly was modestly affected in *Fam210a* cKO hearts compared to control hearts at the early stage (Figure S5G). Then, we examined Δψ_m_ and ROS production in mitochondria of isolated CMs. TMRE staining showed disturbed Δψ_m_ in part of the mitochondria in ∼40% of KO CMs at the early stage (Figure 3D) and an overall increase of Δψ_m_ in *Fam210a* KO CMs (Figure 3E, S5H). This heterogeneity in defects of Δψ_m_ was not observed in control CMs. Moreover, ROS production was significantly increased at the early stage in KO CMs (Figure 3F).

Finally, although we observed a modest overall increase in the intensity of TMRE staining in FAM210A deficient CMs (Figure 3E), seahorse measurement revealed that the mitochondrial respiratory activity was moderately compromised due to disruption of Δψ_m_ in part of the mitochondria (Figure 3G), with a 21.7% decrease of the maximal oxygen consumption rate and a 42.9% decrease of spare respiratory capacity (Figure 3H) in KO CMs compared to control cells at the early stage. Collectively, these results suggest that reduced Δψ_m_ and respiratory activity accompanied by increased ROS production are primary disease driving events, which contribute to the progression of dilated cardiomyopathy in *Fam210a* cKO mice.

### Transcriptomic and proteomic reprogramming occurs at early and late stages of *Fam210a* cKO hearts

We noticed no obvious pathological phenotypes in CMs and the heart of *Fam210a* cKO mice at the early stage (Figure 3A, S4). The cardiac dilation and failure occur progressively during the late stage (Figure 1). To dissect global gene expression changes during HF development and understand the etiology at the molecular level, bulk RNA-seq and quantitative protein mass spectrometry were performed for whole heart tissues harvested from cKO and control mice at ∼5 weeks (early-stage) and ∼9 weeks (late-stage) post KO (Figure 4A, B, Table S2, S3). At the early stage, 124 mRNAs were significantly upregulated, while 49 mRNAs were significantly downregulated in the cKO hearts (Figure 4B, Table S2). In parallel, 169 proteins were increased while 101 proteins were decreased in the early stage cKO hearts (Figure 4B; Table S3). Moreover, no strong correlation was observed between mRNA and protein dysregulated expression at the early stage (Figure 4C). Consistently, only a few genes were overlapped between differentially expressed mRNAs and proteins at the early stage (6 upregulated and 3 downregulated genes) (Figure 4D, Table S4). Fam210a is present as one of the three downregulated genes as a positive control. In contrast, a large number of mRNAs were differentially regulated at the RNA level, with 4370 significantly upregulated genes and 4301 significantly downregulated genes in the late-stage cKO hearts (Figure 4B; Table S2), probably driven by the combined effect of FAM210A deficiency and massive secondary effects of HF at the late stage. In parallel, within detectable proteins by mass spectrometry, 157 proteins were upregulated while 104 proteins were downregulated (Figure 4B; Table S3). Intriguingly, a strong positive correlation (R=0.56) between the expression of mRNAs and proteins was observed at the late stage (Figure 4E), indicating a coordinated regulation of the steady-state mRNA and protein expression at this stage. In agreement with this observation, most of the significantly dysregulated proteins in the proteomic analysis were also significantly dysregulated at the mRNA level. Within overlapped dysregulated genes, 137 genes were upregulated, and 68 genes were downregulated at the late stage (Figure 4F). Notably, the most prominent enriched gene pathways uncovered by gene ontology (GO) analysis in upregulated mRNAs at the early stage include tRNA aminoacylation and translation (Figure 4G, I, Table S4), while downregulated mRNAs were enriched in coagulation and metabolic pathways at the early stage (Figure 4G, Table S4). At the protein level, the upregulated genes were enriched in neutrophil-mediated immunity, while downregulated genes were enriched in mitochondrial translation machinery at the early stage (Figure 4G, I, Table S4). In the overlapped genes at the late stage, tRNA aminoacylation and amino acid metabolism were the most highly enriched pathways in upregulated genes, while sulfide oxidation and mitochondrial ETC assembly were enriched in downregulated genes at the late stage (Figure 4H, J, Table S4).

**Figure 4.**
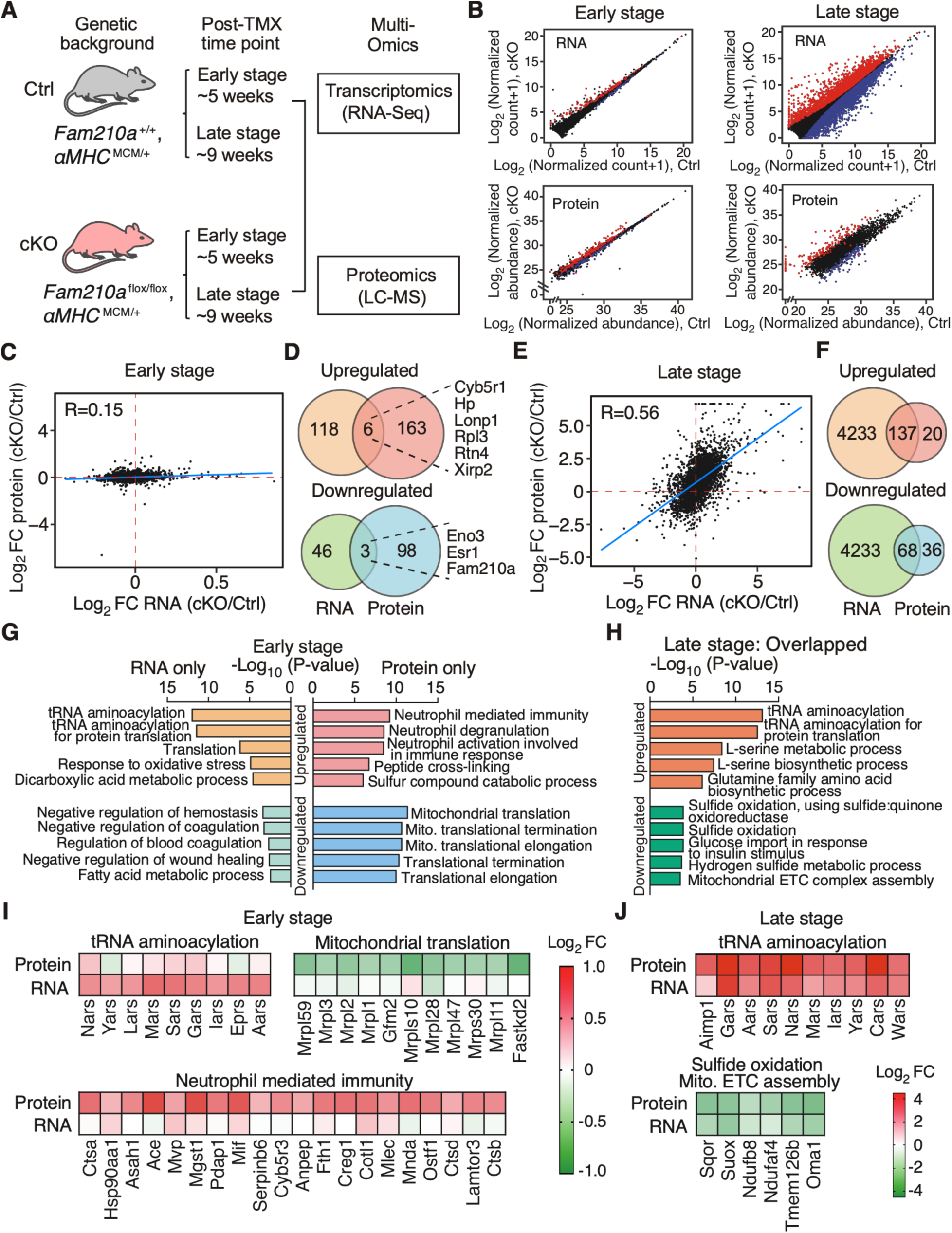
Transcriptomic and proteomic profiles in FAM210A deficient hearts. (**A**) Experimental timeline of sample collection for transcriptomic and proteomic analyses in *Fam210a* cKO hearts. (**B**) Differentially expressed genes at RNA and protein levels in cKO hearts at the early and late stages. Red: significantly increased genes; blue: significantly increased genes. (**C, E**) The Pearson correlation between changes of RNA and protein in cKO hearts at the early (C) and late stages (E). (**D, F**) Venn diagrams illustrate the overlap of dysregulated genes at RNA and protein levels in cKO hearts at the early (D) and late stages (F). (**G**) Gene ontology (GO) analyses of the biological process for dysregulated genes from D at the RNA or protein level in cKO hearts at the early stage. Enrichr web tool was used to conduct the GO analysis. (**H**) GO analyses of biological process for overlapped genes from F at the RNA level in cKO hearts at the late stage. Only the top 5 GO terms were shown in G, H. (**I**) Expression heat map of a selected group of RNA only upregulated genes, protein-only upregulated genes, and protein-only downregulated genes at the early stage from top GO terms of G. (**J**) Expression heat map of select co-regulated genes at the RNA and protein levels at the late stage from top GO terms in H.

To explore the global translation profile upon *Fam210a* cKO *in vivo*, we performed ribosome profiling (Ribo-seq) using sucrose cushion (1M sucrose, 34.2% w/v) ^19, 20^ with whole heart tissue lysates of control and *Fam210a* cKO hearts at the early stage (Figure S5I, S5J, Table S5). This method can capture cytosolic monosomes and high molecular weight mitochondrial monosomes complexed with their ribosome-protected footprints (RPF) upon RNase I digestion. We discovered that 5 mRNAs (*P*_adj_ < 0.05) and 130 mRNAs (*P* < 0.05) had increased RPF density as calculated by Ribo-seq normalized reads divided by RNA-seq normalized reads using the RiboDiff web tool ^21^. Gene ontology results also highlighted the respiratory electron transport chain-related processes, mainly the mitochondrial-encoded genes (MEGs), as a major pathway affected among the top RPF density-upregulated mRNAs (Figure S5K, left). Among these RPF density-increased mRNAs, we noticed multiple MEGs, including *mt-Nd6* (Complex I gene; *P* = 1.17x10^-8^; log_2_FC = 1.41), *mt-Cytb* (Complex III gene; *P* = 1.39x10^-3^; log_2_FC = 0.99), *mt-Co1* (or *Cox1*; Complex IV gene; *P* = 9.30x10^-6^; log_2_FC = 1.04), and *mt-Co3* (or *Cox3*; Complex IV gene; *P* = 0.012; log_2_FC = 1.51). In addition, we observed that *mt-Nd5* (Complex I gene; *P* = 0.040; log_2_FC = 0.50) and *mt-*Nd2 (Complex I gene; *P* = 0.057; log_2_FC = 1.29) show increased RPF density with borderline statistical significance. In contrast, the mRNA expression of these MEGs remained unchanged in the corresponding RNA-seq dataset (Table S5). It is widely accepted that RPF density is inversely proportional to translation efficiency (TE) if translation is regulated at the elongation step ^22^. Thus, FAM210A loss of function may lead to reduced mitochondrial translation elongation and accumulated ribosomes on the MEG mRNAs due to possible ribosome pausing or stalling (Figure S5L). Therefore, these findings support our idea of FAM210A-mediated regulation of translation elongation of MEG mRNAs ^12^ (Figure S5L). On the other hand, we found that 7 mRNAs (*P*_adj_ < 0.05) and 110 mRNAs (*P* < 0.05) showed decreased RPF density in our Ribo-seq data (Figure S5J, Table S5). All of these mRNAs, enriched in multiple gene ontology terms related to regulation of JNK cascade, elastic fiber assembly, protein ubiquitination, and TORC1 signaling, are encoded by the nuclear genome and translated inside the cytoplasm by cytosolic ribosomes (Figure S5K, right). It is well known that RPF density is proportional to TE if translation is regulated at the initiation step ^22^ (Figure S5L). These results suggest a limited number of mRNAs are downregulated at the translation initiation step as a secondary effect following *Fam210a* cKO at the early stage. This secondary effect is likely resulting from integrated stress response, a common signature of mitochondrial cardiomyopathy, driven by phosphorylation and inactivation of eukaryotic translation initiation factor 2α (eIF2α) ^23^.

### Integrated stress response is persistently activated in *Fam210a* deficient hearts

Since RNA-seq analysis provides much deeper genome-wide information about adaptive response and gene regulation than proteomics, we sought to determine the dysregulated gene signature from early and late stages (Figure 5A, Table S2). Overlapping significantly dysregulated genes from RNA-seq datasets revealed 95 upregulated genes and 29 downregulated genes at both stages (Figure 5B, S6A, Table S6). In comparison, a large number of unique dysregulated genes (4275 upregulated and 4272 downregulated genes) were uncovered at the late stage compared to a unique cohort of 29 upregulated and 21 downregulated genes at the early stage (Figure 5B, S6A, Table S6). GO analysis of overlapped upregulated genes identified that aminoacyl-tRNA synthetases, amino acid synthases, and amino acid transporters were enriched, indicating a persistent activation of the integrated stress response (ISR) in *Fam210a* cKO hearts at both stages (Figure 5C, D, Table S6). Meanwhile, GO analysis of unique dysregulated mRNAs at the late stage showed that proteasome and ribosome pathways were upregulated (Figure S6B, C, Table S6), suggesting a futile cycle of increased protein degradation and synthesis machinery, which may maintain the proteostasis in KO CMs as a compensatory response. On the other hand, the ETC pathway was downregulated (Figure S6B, D). We also observed decreased expression of most fatty acid β-oxidation related metabolic enzymes and increased expression of more than half the glycolysis related metabolic enzymes (Figure S6E), which is consistent with the fatty acid-to-glycolysis metabolic switch during HF.

**Figure 5.**
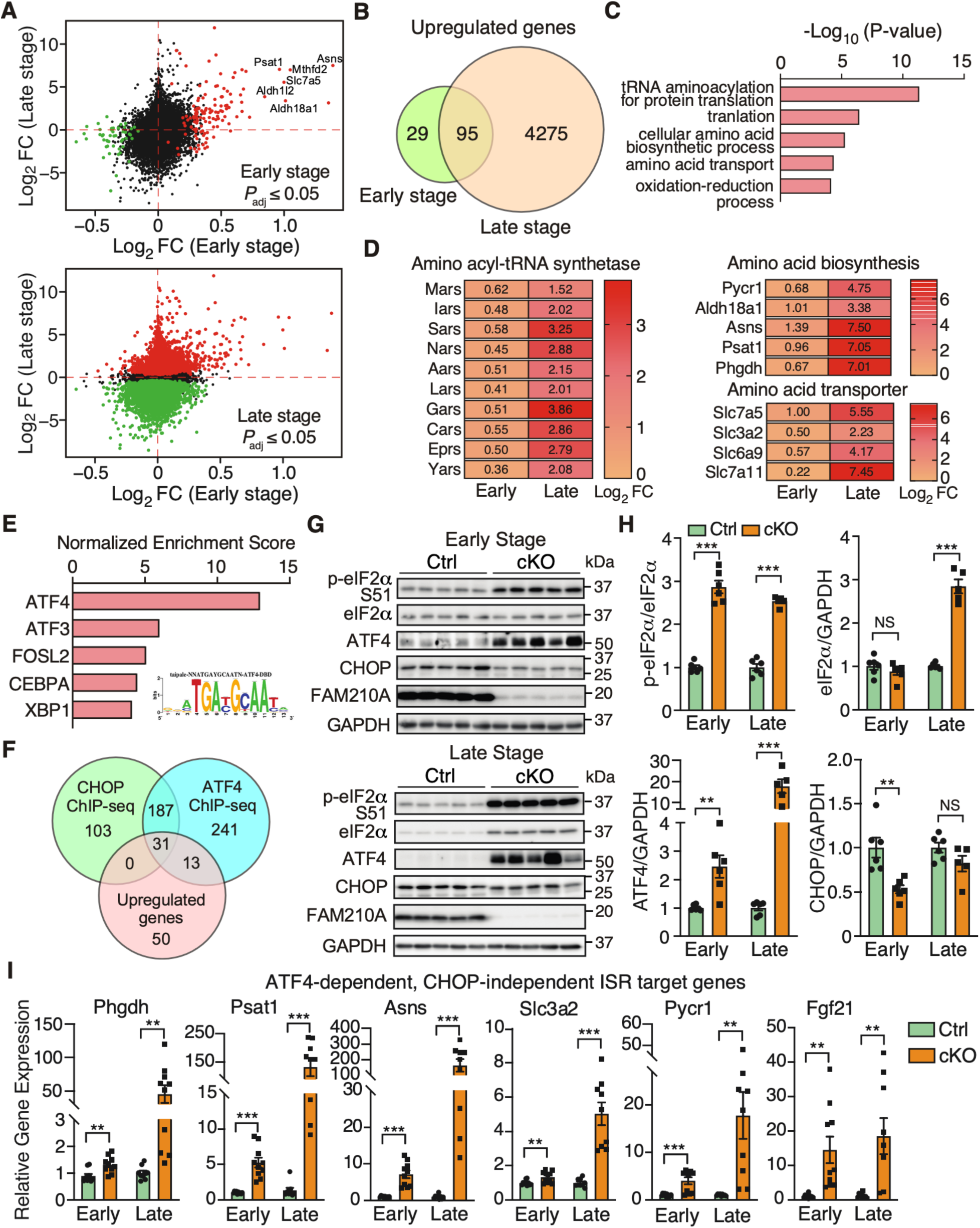
Cardiomyocyte-specific FAM210A deficiency leads to persistent activation of ATF4-dependent CHOP-independent integrated stress response. **(A)** Differentially expressed genes at the transcriptomic level at the early (top) and late stages (bottom). (**B**) The Venn diagram illustrates the overlap of upregulated genes at the RNA level in *Fam210a* cKO hearts at the early and late stages. (**C**) GO analysis using the Enrichr web tool suggests enhanced amino acid metabolism-related gene expression in cKO hearts. (**D**) Expression heat map of the differentially expressed genes from top 3 GO terms in C. (**E**) iRegulon analyses identified ATF4 as the top transcription factor with the most enriched regulatory motif in all the 95 genes (from B) in the inset. (**F**) Overlapping upregulated genes with CHOP and ATF4 ChIP-Seq databases reveals ATF4 regulated target genes in CMs. (**G**) Western blot detection of phospho-eIF2α, ATF4, and CHOP at the early and late stages of cKO hearts. (**H**) Quantification of protein expression as shown in G. N = 6 for Ctrl and cKO at the early stage. N = 6 for Ctrl and N = 5 for cKO at the late stage. (**I**) Relative RNA expression of ATF4-dependent, CHOP-independent ISR downstream targets in cKO hearts at the early and late stages. β-actin was used as a normalizer. N = 5M + 5F for Ctrl and cKO at the early stage. N = 5M + 4F for Ctrl and cKO at the late stage. NS, not significant; ** *P* < 0.01; *** *P* < 0.001 by student t-test (H, I).

Consistent with the activation of ISR, transcription factor analysis of overlapped upregulated genes using the iRegulon tool ^24^ revealed that activating transcription factor 4 (ATF4) was the dominant transcription factor potentially involved in the transcriptional activation of the ISR target genes (Figure 5E). Activation of ISR often induces the expression of ATF4 and CCAAT-enhancer-binding protein homologous protein (CHOP), which is responsible for initiating cell apoptosis ^25^. To distinguish the transcriptional effects from ATF4 or CHOP, we overlapped persistently upregulated genes with existing CHOP and ATF4 ChIP-Seq data ^26^. We found that some ISR targets were unique for ATF4, while no CHOP-specific targets were found (Figure 5F, Table S7). Western blot analysis confirmed activation of ISR as indicated by phosphorylation of translation initiation factor eIF2α and increased protein and mRNA expression of ATF4 (Figure 5G, H, S7A). Intriguingly, the CHOP protein level was reduced at the early stage (Figure 5G, H), suggesting a CM-specific ATF4-driven, CHOP-independent ISR program in the cKO hearts. We confirmed that the CHOP independent, ATF4 target genes were significantly increased in KO CMs at both stages, including phosphoglycerate dehydrogenase (Phgdh), phosphoserine aminotransferase 1 (Psat1), asparagine synthetase (Asns), solute carrier family 3 member 2 (Slc3a2), pyrroline-5-carboxylate reductase 1 (Pycr1), and fibroblast growth factor 21 (Fgf21) (Figure 5I). In addition, the shared target genes of ATF4 and CHOP were also remarkably increased at both stages, including activating transcription factor 5 (Atf5), solute carrier family 7 member 5 (Slc7a5), phosphoserine phosphatase (Psph), methylenetetrahydrofolate dehydrogenase (NADP^+^ dependent) 2/ methenyltetrahydrofolate cyclohydrolase (Mthfd2), aldehyde dehydrogenase 1 family member L2 (Aldh1l2), aldehyde dehydrogenase 18 family member A1 (Aldh18a1), serine hydroxymethyltransferase 2 (Shmt2), and growth differentiation factor 15 (Gdf15) (Figure S7A). In contrast, we only observed an increase of mRNA of three mitochondrial unfolded protein response (UPR^mt^) marker genes at the late stage, including heat shock protein family D member 1 (Hsp60), heat shock protein family A member 4 (Hsp70), and mitochondrial lon peptidase 1 (LonP1) (Figure S7B). Only LONP1 protein expression was increased at the late stage (Figure S7C, D), suggesting that UPR^mt^ is not a major downstream effector pathway caused by FAM210A deficiency at the early stage in the heart. Four kinases are known to phosphorylate eIF2α and activate the ISR upon various stresses ^27^. In FAM210A deficient hearts, we only observed a band upshift of Heme-Regulated Inhibitor (HRI) kinase while the other three kinases (GCN2, PEKR, and PKR) were unchanged in their phosphorylation at the early stage (Figure S7E, F), suggesting that HRI may be the kinase activating the ISR caused by *Fam210a* deficiency and mitochondrial translation dysregulation. This observation supports the idea that HRI kinase senses mitochondrial damage ^28^ and activates the ISR upon mitochondrial stresses in the heart. However, all four kinases were activated at the late stage, probably due to various stresses in HF. These results suggest that chronic ISR is activated in *Fam210a* KO CMs, which may contribute to the etiology of HF.

### Early-stage metabolomic alterations are present before cardiac dysfunction in *Fam210a* cKO hearts

Our transcriptomic and proteomic analyses demonstrate that a persistent ISR is activated as indicated by the upregulation of multiple ISR downstream target genes involved in amino acid biosynthesis (Asns, Aldh18a1, Pycr1, Phgdh, Psat1), nucleotide synthesis (Ctps, Pck1), and amino acid transport (Slc38a1, Slc3a2, Slc6a9, Slc7a11, Slc7a5) in the *Fam210a* cKO hearts (Figure 5, Table S2). Therefore, we ask whether transcriptomic and proteomic reprogramming of these metabolism related pathways in *Fam210a* cKO hearts may remodel the overall metabolomic profile. To address this question, we used liquid chromatography-tandem mass spectrometry (LC-MS/MS) to evaluate the metabolomic profile of the hearts of cKO and control mice. Principal component analysis suggests that the overall metabolic profile in cKO hearts (red) is different from that in controls (green) at the early stage of MC (30-day post-KO) (Figure S8A, Table S8) as well as at the late stage (60-day post-KO) (Figure S8B, Table S8). Consistent with the alteration in ISR downstream metabolic enzymes at the early stage (Figure 5A, D), Metabolite Set Enrichment Analysis (Figure S8C) and top dysregulated metabolites in heat maps revealed that amino acid and nucleotide biosynthetic pathways are among the top changed metabolic pathways, including multiple dysregulated amino acids and nucleotide precursors (Figure S8D). These metabolic pathways stayed dysregulated throughout the late stage of cKO hearts (Figure S8E-F), such as reduced histidine and glutamine, increased phenylalanine and ornithine, and increased nucleotide synthesis (Figure S8G). Additionally, several other metabolic pathways were also remarkably changed at the late stage (Figure S8E, F), probably due to massive gene dysregulation at the mRNA and protein levels caused by HF (Figure 5A), such as the glycolysis pathway (Figure S6E), arachidonic acid metabolism, and estrogen metabolism. Interestingly, we observed a persistent increase of glutathione and glutathione disulfate at early and late stages (Figure S8G), indicating a compensatory response to maintain the cellular redox balance in response to enhanced mitochondrial ROS production in cKO hearts. Collectively, these metabolomic analyses indicate that FAM210A deficiency in CMs remodels glycolysis and citric acid cycle-centered amino acids and nucleotide metabolism in the heart, which contributes to the pathogenesis underlying cardiomyopathy and HF.

### FAM210A depletion reduces mitochondrial-encoded mRNA translation in murine hearts

Our prior studies indicate that FAM210A interacts with mitochondrial translation elongation factor EF-Tu and positively regulates mitochondrial-encoded protein expression ^12^. However, whether FAM210A regulates mitochondrial-encoded proteins at the translational level in the heart *in vivo* is unclear. Considering mitochondrial translational defects as a common trigger of ISR ^29–31^, we sought to examine any potential mitochondrial translation defects in the FAM210A deficient hearts. To prove that loss of FAM210A influences mitochondrial translation, we performed mitochondrial polysome profiling using 10-30% sucrose gradient centrifugation coupled with RT-qPCR in isolated mitochondria from the early stage *Fam210a* cKO and control hearts (Figure 6A, B). Translation efficiency of mitochondrial-encoded gene mRNAs was significantly reduced in cKO hearts compared to controls, as indicated by decreased mRNA association with translationally active fractions of mitochondrial-encoded genes (Figure 6C-G). Consistently, we observed moderate but significant reductions at the steady state levels in multiple mitochondrial-encoded ETC proteins in isolated mitochondria from *Fam210a* cKO hearts compared to control hearts at the early stage, including ND1, CYB, COX2, ATP6 using global protein loading as a normalizer (Figure 6H, I). In contrast, we did not observe significant changes in the nuclear-encoded mitochondrial proteins (NDUFS1, SDHA, UQCRC2, ATP5B) except COX4. Considering the role of regulating mitochondrial protein expression by FAM210A ^12^, we reason that a slightly increased mitochondrial DNA copy number (Figure 3C) is unable to compensate for any reduced translation efficiency of mitochondrial-encoded proteins caused by loss of FAM210A. Consistent with this idea, when mitochondrial DNA copy number was reduced at the late stage (Figure 2H), we observed that the steady-state level of mitochondrial-encoded proteins and nuclear encoded proteins was significantly reduced in *Fam210a* cKO hearts, as shown by mass spectrometry (Figure S9A, Table S3) and confirmed by Western blot analyses (Figure S9B, C). Taken together, these results suggest that loss of FAM210A compromises the translation of multiple mitochondrial-encoded ETC component genes *in vivo* as the trigger for initiating mitochondrial dysfunction.

**Figure 6.**
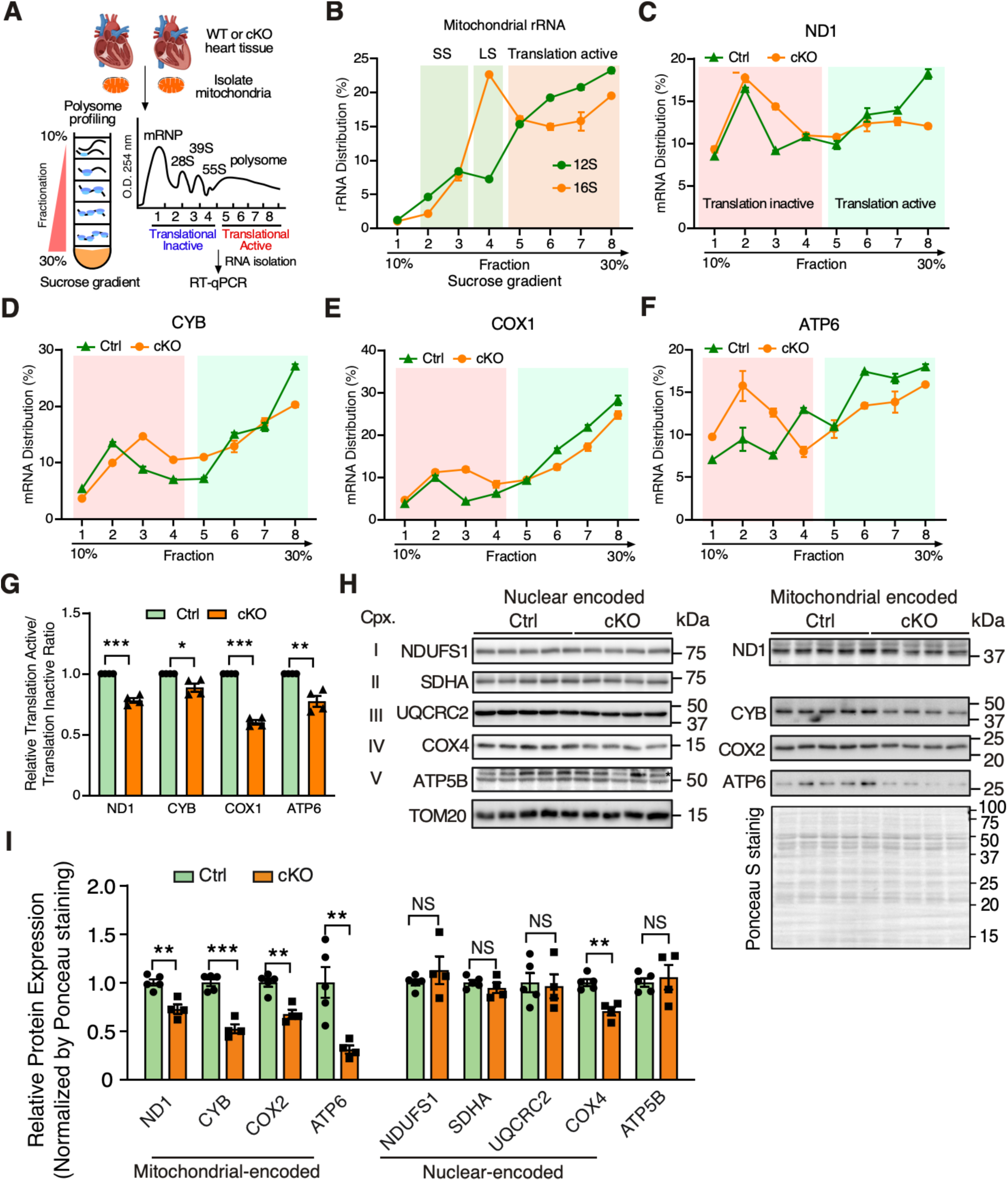
FAM210A regulates mitochondrial translation in the heart. (**A**) Schematic of the workflow of polysome profiling analysis of cardiac mitochondrial lysates from WT and cKO mouse hearts. (**B**) RT-qPCR detection of mitochondrial 12S and 16S rRNA. SS: small mitoribosome subunit; LS: large mitoribosome subunit (**C-G**) RT-qPCR analysis of mitochondrial-encoded mRNA distributions of *Nd1*, *Cyb*, *Cox1*, and *Atp6* across different mitochondrial polysome profiling fractions (C-F). The quantification of the mRNA distribution ratio between translation active and inactive fractions was shown in G (N = 4 biological replicates). (**H, I**) Western blot measurement of nuclear and mitochondrial-encoded ETC protein levels using isolated mitochondria from early stage Ctrl or *Fam210a* cKO hearts at 5-week post gene knockout (H). The quantification results are shown in I. N = 5 for Ctrl and N = 4 for cKO. * *P* < 0.05; ** *P* < 0.01; *** *P* < 0.001 by student t-test (G, I).

### FAM210A overexpression alleviates cardiac pathological remodeling in myocardial infarction

Compromised mitochondrial activity is a common feature in hearts under ischemic stresses, and an increase in mitochondrial biogenesis is considered a promising therapeutic strategy to treat ischemic heart disease^7^. FAM210A was previously reported to be reduced by 34% in murine hearts at the mRNA level under MI versus sham surgical conditions by a microarray-based transcriptome screening ^32^. After examining the expression of FAM210A in human HF samples, we found that FAM210A protein expression was significantly decreased in the hearts of ischemic HF (IHF) patients compared to non-failing human hearts (Figure 7A). However, *FAM210A* mRNA showed a reduction trend without statistical significance (Figure S9D). Moreover, the FAM210A protein level was also reduced in the mouse MI hearts compared to sham-controlled ones (Figure 7B). The mRNA level was significantly reduced in the MI compared to sham hearts (Figure S9E). In addition, we observed reduced expression of mitochondrial-encoded proteins in human IHF and mouse MI heart tissue samples compared to their corresponding non-failing controls (Figure S9F, G). As we uncovered the essential role of FAM210A in maintaining mitochondrial mRNA translation (Figure 6, S9) and observed reduced FAM210A protein expression in human and mouse IHF samples, we sought to determine whether overexpression of FAM210A would enhance the mitochondrial protein expression and reduce cardiac damage using an MI mouse model (Figure 7C). To test this hypothesis, we subcutaneously injected control GFP and FAM210A-overexpressing adeno-associated viruses (AAV9) in WT mice at around postnatal day P5. MI surgery was performed in *Fam210a* AAV9 overexpression (OE) and control mice at ∼8 weeks followed by left anterior descending artery (LAD) ligation for 4 weeks, and the cardiac function was monitored biweekly by echocardiography. At the end-point, FAM210A OE was confirmed by immunoblot in the hearts at 4 weeks post LAD ligation (Figure 7D), and the cardiac function was significantly improved in MI mice with FAM210A OE compared to the control AAV treatment (Figure 7E, Table S9). The HW/TL ratio of MI hearts with FAM210A OE was decreased by 16.6% (Figure 7F). Moreover, the scar area was reduced by FAM210A OE, as indicated by Picrosirius red staining (Figure 7G). We quantified infarct size normalized to area at risk (AAR) using an echocardiographic analysis. We did not notice any difference in AAR while the infarct size was significantly decreased after normalization by AAR (Figure 7H). Consistently, CM apoptosis was also significantly reduced by 57.9% at the border zone, as indicated by TUNEL staining (Figure 7I). Western blot analysis showed that mitochondrial-encoded protein expression was enhanced by FAM210A overexpression, including ND1, CYB, COX2, and ATP6 (Figure 7J). In contrast, nuclear-encoded mitochondrial protein expression was not affected except COX4, indicating a translation regulatory function of FAM210A in promoting mitochondrial-encoded protein expression *in vivo*. Moreover, ATP production and mitochondrial respiratory activity were increased in FAM210A OE hearts (Figure S10A, B), suggesting an improved bioenergetics and mitochondrial function in MI hearts. As a result, ISR activation was compromised after FAM210A OE in MI mice, as indicated by reduced p-eIF2α/eIF2α ratio at the protein level (Figure S10C). Taken together, these data suggest that FAM210A overexpression protects hearts from cardiac damage and functional decline under ischemic stress.

**Figure 7.**
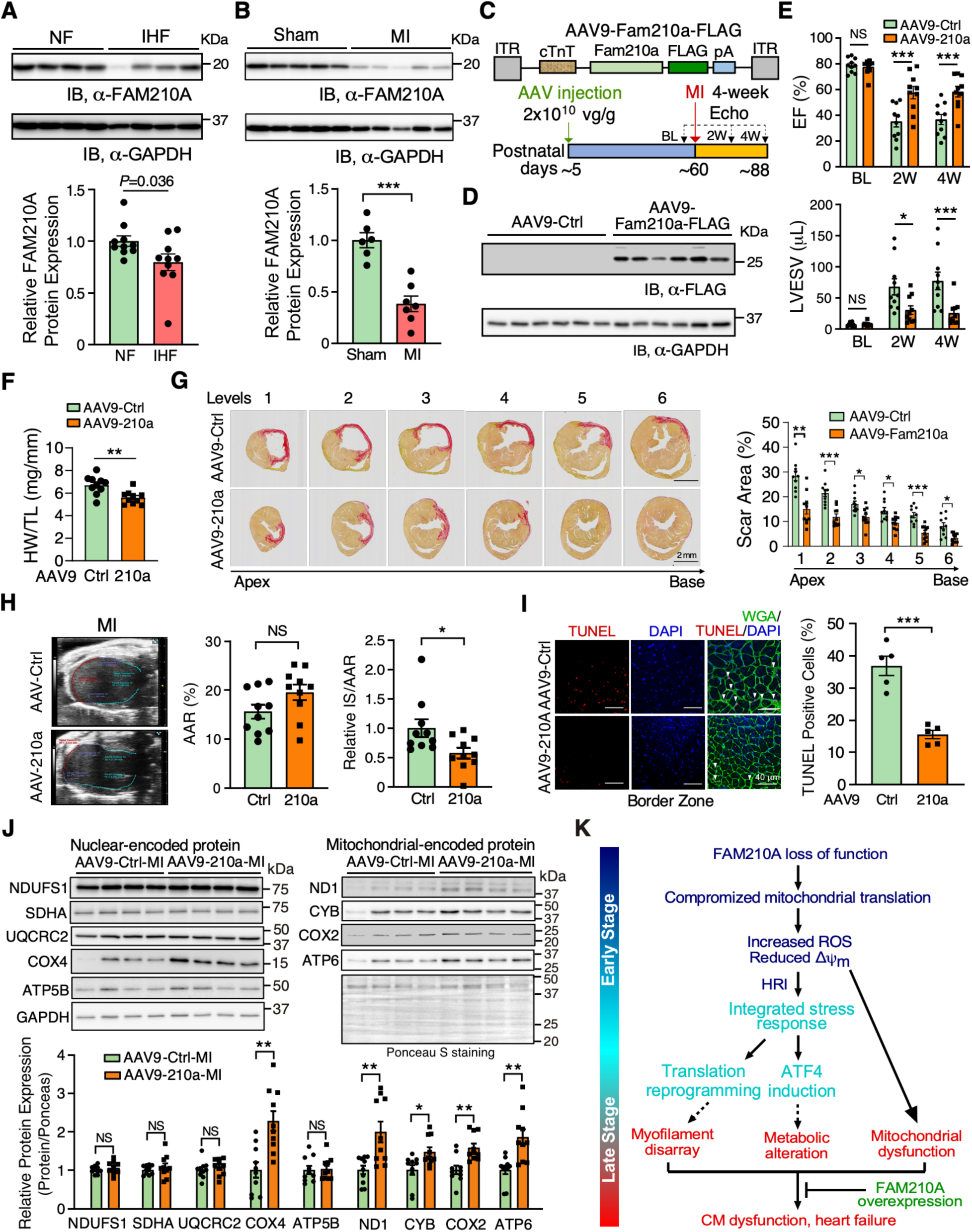
FAM210A overexpression improved cardiac function after myocardial infarction. (**A**) FAM210A protein expression in human ischemic heart failure (IHF) vs. non-failing (NF) hearts. N = 10 for NF; N = 10 for IHF. (**B**) FAM210A protein expression in murine hearts from MI (2 weeks) vs. sham-treated mice. N = 6 for sham; N = 7 for MI. (**C**) Workflow of the AAV9-FAM210A-FLAG overexpression in an MI model. AAV9-GFP was used as a negative control treatment. (**D**) Western blot detection of FAM210A protein at 4-week post-MI. (**E**) Left ventricular function was measured by echocardiography in *Fam210a* overexpression and control mice with MI surgery. EF: ejection fraction; LVESV: left ventricle end-systolic volume; LVEDV: left ventricle end-diastolic volume. N = 10 for Ctrl (6M + 4F) and overexpression (3M + 7F). (**F**) HW/TL ratio in *Fam210a* overexpression mice. N = 10 for Ctrl (6M + 4F) and overexpression (3M + 7F). (**G**) Picrosirius red staining of the scar area in the hearts of Ctrl and *Fam210a* overexpression mice at 4 weeks post-MI. Right panel: quantification of the scar area at each level. (**H**) Measurement of AAR and infarct areas using echocardiographic analysis. N = 10 for AAV-Ctrl and AAV-210a. NS, not significant. AAR: area at risk; IS: infarct size. (**I**) Representative images of CM apoptosis in hearts of mice subjected to MI operation. Red color indicates TUNEL positive (apoptotic) nuclei, green indicates CM staining by WGA, and blue (DAPI) represents nuclei. The percentile of TUNEL-positive nuclei was quantified in the right panel. (**J**) Western blot analysis of nuclear-encoded mitochondrial ETC proteins and mitochondrial-encoded ETC proteins. (**K**) Schematic working model of the impact and consequence of FAM210A loss-of-function (cKO) and gain-of-function (AAV9 OE) in the heart. * *P* < 0.05; ** *P* < 0.01; *** *P* < 0.001 by two-way ANOVA with Sidak multiple comparisons test (E, G), student t-test (B, F, H, I, J), and Mann-Whitney test (A).

## Discussion

This study demonstrates a novel molecular function of FAM210A in mitochondria, CMs, and the heart using genetic knockout mouse models. FAM210A regulates mitochondrial mRNA translation, and loss of FAM210A causes reduced mitochondrial protein synthesis. The mitochondrial translational defect causes disrupted Δλφλ_m_ and increased ROS that activates ISR and subsequent remodeling of the cellular transcriptome, translatome. proteome, and metabolism in CMs, leading to dysfunction of CM contraction, eventually causing HF and organismal mortality. We also observed reduced FAM210A protein expression in IHF patients’ whole hearts and MI mouse hearts. The overexpression of FAM210A in mouse CMs enhances mitochondrial protein expression and protects the heart from MI-induced cardiac damage and dysfunction. These results suggest that FAM210A is a novel mitochondrial translation regulator to maintain mitochondrial homeostasis in cardiomyocytes and offers a new therapeutic target for cardiac disease (Figure 7K).

### Translation regulatory function of FAM210A in cardiac mitochondria *in vivo*

Our prior *in vitro* cell culture studies show that FAM210A regulates mitochondrial-encoded protein expression, possibly through interacting with EF-Tu and mitoribosome, anchoring the mitochondrial translation machinery in the inner mitochondrial membrane ^12^. To our knowledge, the current study provides the first evidence to support the idea that FAM210A influences mitochondrial translation *in vivo* using the CM-specific cKO mouse model and AAV9-mediated CM-specific overexpression in the MI mouse model. *αMHC*^MCM^ and *Fam210a* fl/fl mice were used as controls to rigorously support our phenotypic characterizations (Figure 1D, S1G-J). Our Ribo-seq and polysome profiling data show that mitochondrial translation was decreased in FAM210A deficient CMs at the early stage before HF (Figure S5I-L, Figure 6A-G). Consistently, the steady-state level of mitochondrial-encoded proteins was moderately and significantly reduced at this early stage without apparent changes in nuclear-encoded mitochondrial proteins (Figure 6H, I). However, this proteomic imbalance status was worsened with time, as indicated by the significant decrease of mitochondrial-encoded proteins and even nuclear-encoded mitochondrial proteins at the late stage as cardiac dysfunction occurs at ∼9 weeks post-TMX-induced *Fam210a* KO (Figure S9A-C). Thus, the primary events in mitochondria before cardiac dysfunction are likely the disease-driving factors, including disrupted Δλφλ_m_, increased ROS production, and decreased respiratory rate (Figure 3). Before these mitochondrial defects, the initial trigger of such early events is the loss of FAM210A, which leads to an aberrant mitochondrial translation (Figure 7K). These findings imply that reduced FAM210A in IHF patients’ hearts or MI mouse hearts may contribute to compromised mitochondrial translation, disrupted proteostasis, and mitochondrial dysfunction. Given the fact that FAM210A and mitoribosome proteins were the top binding partners of mitochondrial translation-related protein ATAD3A ^17^ and EF-Tu ^12^, we speculate that FAM210A may form a complex with ATAD3A, which acts as a mitochondrial inner membrane-associated scaffold or adaptor to increase the local concentration of EF Tu as well as mitoribosomes to enhance translation efficiency at the elongation step ^12^. *Fam210a* cKO in CMs may reduce EF-Tu docked on the mitochondrial inner membrane, leading to impaired channeling of aminoacyl-tRNA to the mitoribosome decoding center. This slows down the translocation and movement of mitoribosome (reduced elongation rate) and accumulates more mitoribosomes on mitochondrial-encoded mRNAs, thereby increasing RPF density (more ribosome footprints) (Figure S5I-L). Consequently, *Fam210a* cKO activates the integrated stress response that inhibits cytosolic translation initiation due to p-eIF2α, leading to reduced loading of cytoribosome on cytoplasmic mRNAs, thereby decreasing RPF density (fewer ribosome footprints) (Figure S5I-L). Interestingly, we also observed dysregulated cristae structure in *Fam210a* cKO hearts at the early stage, which further supported its functional association with ATAD3A ^17, 18^. Loss of FAM210A did not alter the expression of ATAD3A at the early stage (Figure S5B), indicating that the FAM210A-ATAD3A complex may exert dual functions in mitochondria, including recruiting EF-Tu and mitoribosome for mitochondrial translation, and regulating the mitochondrial cristae structure. Further experiments are required to elucidate the detailed molecular and functional connections between FAM210A with ATAD3A. The FAM210A gene is conserved in more than 213 organisms, including humans, mice, zebrafish, and *C.elegans* ^33^. *Fam210a* mRNA is expressed in all organs and is enriched in the heart and testis of humans ^34^. Therefore, the biological function and molecular mechanism we discover here may be generalizable, and further studies are required to prove FAM210A function across multiple cell types and other species.

### ISR contributes to disease progression of mitochondrial cardiomyopathy

The mammalian target of rapamycin (mTOR) pathway is often associated with cardiac pathological hypertrophy ^35^. We measured the phosphorylation of mTOR kinase and its two substrate proteins, P70S6K and 4EBP1, and found *Fam210a* KO activated neither (data not shown). Moreover, we did not observe a strong UPR^mt^ activation. At the same time, ISR was markedly activated in *Fam210a* cKO hearts at the early stage (Figure 5G, H, S7E, F), suggesting that ISR is a central pathway activated by FAM210A deficiency. ISR is a major translational control pathway to promote the survival of cells under stress conditions in the acute phase and can lead to cell apoptosis after chronic activation ^27^. Our multi-omics results reveal that ISR is persistently activated at early and late stages in *Fam210a* cKO hearts (Figure 7K). Intriguingly, we observed limited but significantly increased (∼2.5%) CM cell death at the late stage of *Fam210a* cKO hearts (Figure S3F), as indicated by TUNEL (terminal deoxynucleotidyl transferase dUTP nick end labeling) staining. This may cause chamber dilation over time and explains the slightly increased cardiac fibrosis at the late stage. However, as downstream effectors of ISR, only ATF4 but not CHOP was upregulated at the protein level (Figure 5G, H). ATF4 activates pro-survival gene expression without triggering a pro-apoptosis gene program mainly activated by CHOP ^27^. Therefore, this ISR seems to be anti-apoptotic for a prolonged time to minimize CM death, as indicated by increased expression of an anti-apoptotic marker gene Mcl1 and no drastic change in pro-apoptotic genes such as Bcl2l11 and Apaf1 (Figure S10D). This “double-edged sword” effect of ISR activated in *Fam210a* cKO hearts can partially explain the delayed onset of severe cardiac pathological remodeling and heart failure (∼7 weeks post-TMX treatment) in *Fam210a* cKO hearts in addition to the relatively long half-life of FAM210A protein ^36^. However, the ISR-mediated adaptive translational response may protect against cell death at the expense of disrupted proteomic homeostasis, leading to aberrant sarcomeric protein expression and contractile dysfunction in CMs (Figures 2 and 4). We observed that robust activation of two effectors reprograms the cellular transcriptome, translatome, proteome, and metabolism, in response to stresses in *Fam210a* cKO hearts, including activated phosphorylation of eIF2α and increased protein expression of ATF4. Phosphorylated eIF2α inhibits cap dependent mRNA translation ^37^. This translational defect inhibits the synthesis of myofilament proteins required for contractile function and nuclear-encoded mitochondrial proteins (Figure S2D, S9A), which causes CM contractile dysfunction. On the other hand, increased ATF4 activates the transcription of multiple metabolic enzymes (Figure 5). As a consequence, amino acid and nucleotide metabolism is altered, and this change may contribute to the progression of HF. Thus, it is worthwhile to test if targeting the ISR pathway provides any beneficial effect in treating MC and MI based on the fact that genetic deletion of ISR kinase PKR protects hearts from pressure overload-induced HF ^38^ and that chemical inhibition of ISR by ISRIB prevents atrial fibrillation post-MI ^39^. Our integrated Ribo-seq and RNA-seq analysis in both cKO and control *αMHC*^MCM^ hearts at the early stage supports the role of FAM210A in regulating mitochondrial mRNA translation elongation *in vivo* (Figure S5I-L). It is well known that compromised cytosolic translation elongation triggers ISR activation ^40, 41^. Here, we provide evidence to support the notion that impaired mitochondrial translation elongation (Figure S5I-L) can also activate the ISR pathway possibly via HRI kinase (Figure S7E, F). Heme or iron deficiency enhances phosphorylation of eIF2α through HRI and activates ISR, thereby affecting cytosolic translation ^42^. The ISR activation affects the synthesis of mitochondrial ribosomal proteins, which are translated in cytosol, thereby regulating the protein synthesis inside mitochondria during erythropoiesis ^42^. In our study, HRI is activated by mitochondrial translation deficiency at the early sage in *Fam210a* cKO hearts. Similar to the effect in erythroblasts, HRI may indirectly influence mitochondrial translation via a secondary effect to some extent in *Fam210a* deficient CMs, as indicated by enriched mitochondrial translation elongation and termination factors in downregulated proteins in our early-stage mass spectrometry data (Figure 4G).

### FAM210A overexpression protects hearts during myocardial infarction

In contrast to reduced mitochondrial-encoded mRNA translation by genetic knockout of *Fam210a* in murine hearts, we show that overexpression of FAM210A positively regulates mitochondrial translation and protects hearts from pathological remodeling triggered by ischemic stress during MI (Figure 7). We assume that reduced FAM210A expression in human IHF and mouse MI hearts may contribute to a compromised compensatory boost in mitochondrial mRNA translation and protein synthesis in the mitochondrial matrix ^43^, disrupts mitochondrial homeostasis, and exaggerate downstream mitochondrial and CM dysfunction. Therefore, overexpression of FAM210A can enhance mitochondrial translation in CMs, improve mitochondrial respiratory activity and energetics (ATP production), and protect hearts under ischemic stresses by promoting mitochondrial activities (Figure S10A-C, 7K). This phenotypic rescue by overexpression of FAM210A is consistent with the proposed mechanism and benefits of maintaining a balance of cytosolic and mitochondrial translatomes and translation programs in yeast and human cells ^44, 45^. A prior multi-omics study in animal models shows an increased FAM210A expression in swim-activated physiological hypertrophy compared to pressure overload-induced cardiac pathological hypertrophy ^14^, supporting the potential cardiac protective function of FAM210A overexpression. Further studies are required to examine whether FAM210A overexpression shows any cardiac protection in non-ischemic HF models since we observed exaggerated cardiac pathological hypertrophy in a pressure overload HF mouse model with induced FAM210A expression upon genetic ablation of miR-574 ^12^.

## Data availability

RNA-seq data produced and used in this study were deposited on Gene Expression Omnibus (GEO) database under accession number GSE195957.

## Funding

This work was supported in part by National Institutes of Health grants R01 HL132899, R01 HL147954, and R01 HL164584 (to P.Y.), start-up funds from Aab Cardiovascular Research Institute of the University of Rochester Medical Center (to P.Y.), American Heart Association Postdoctoral Fellowship 19POST34400013 and Career Development Award 848985 (to J.W.), NIH T32 Fellowship (T32 GM068411 to O.H.), and National Institutes of Health grant R01 HL154318 (to C.Y.).

## Acknowledgments

We are grateful to Jared Hollinger, Eng-Soon Khor (Aab CVRI), and Bin Liu (Mississippi State University) for their critical reading of the manuscript and Qiuqing Wang (Cleveland Clinic) for biostatistical consulting. We appreciate the technical assistance from Erika Flores Medina, Mohan Amy, and Deanne Mickelsen (Aab CVRI) in histology, surgical, and echocardiography operations (AAR image analysis), respectively. RNA seq and primary data analysis were performed by Jason R Myers from the Genomics Research Center at the University of Rochester. Genomics Research Center at the University of Rochester and Novogene company performed Ribo-seq (RNA-seq) and data analysis. We appreciate the bioinformatic data analysis for Ribo-seq and RNA-seq performed by Dalia Ghoneim. Kevin Welle performed protein mass spectrometry analysis from the Mass Spectrometry Resource Lab at the University of Rochester. Chad Galloway and Karen de Mesy Bentley conducted the electron microscopic imaging of heart sections in the Electron Microscopy Core at the University of Rochester. Eric Small provided us with the tamoxifen-inducible transgenic mouse line *αMHC*^MerCreMer^ (*αMHC*^MCM/+^). Hani Awad and Emma Gira contributed to testing the potential changes in bone morphology and function in *Fam210a* cKO mice. None of the authors have any financial conflict of interest related to the research described in this manuscript.

## Conflict of Interest Disclosures

None.

## Author Contribution Statement

PY obtained the funding and launched the study. PY and JW conceived the ideas, designed the experiments, analyzed the data, and wrote the manuscript. JW and KCVS carried out the experimental work. OH and SC provided technical assistance and performed specific experiments. JM, WHWT, and CY provided conceptual feedback, technical support, and research platforms for metabolomic analysis, human sample examination, and cardiomyocyte functional characterization, respectively. All the authors discussed the results and had the opportunity to comment on the manuscript.

## Supplementary Information for

### Supplemental figures

**Figure S1.**
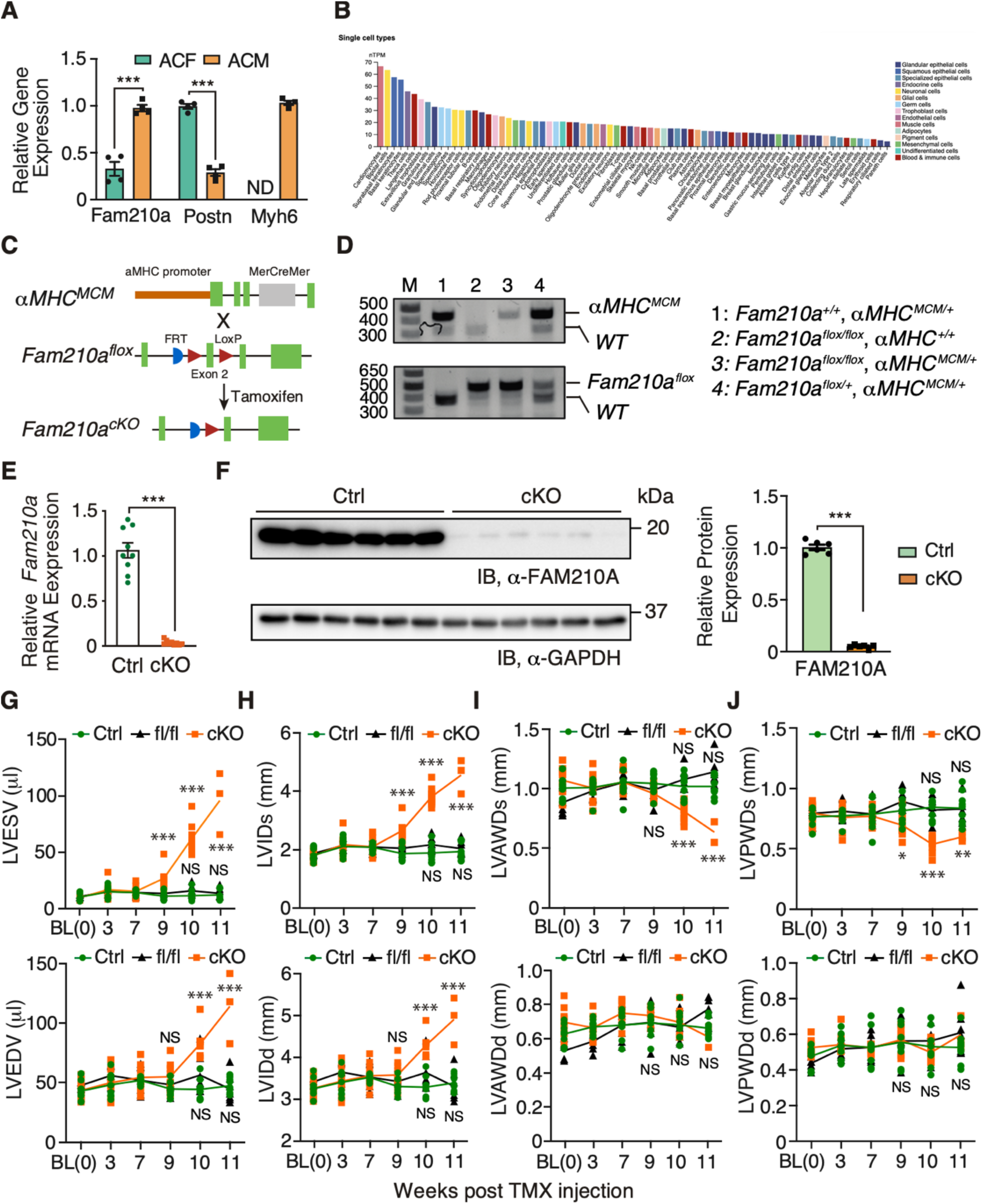
Generation of cardiomyocyte-specific *Fam210a* conditional knockout mice. (**A**) *Fam210a* mRNA expression in isolated adult cardiomyocytes and cardiac fibroblasts. (**B**) *FAM210A* mRNA expression in different cell types across all human organs from single-cell sequencing results of Human Protein Atlas. (**C**) The strategy of generating tamoxifen-inducible CM-specific *Fam210a* conditional knockout mice. (**D**) Genotyping of *Fam210a* cKO and control mice. (**E, F**) FAM210A mRNA (E) and protein (F) expression in the hearts of control and cKO mice ∼65 days post tamoxifen administration. (**G-J**) Cardiac chamber geometric parameters were measured by echocardiography. N = 5M + 2F for *Fam210a* fl/fl control mice. NS: not significant for fl/fl versus Ctrl by Two-way ANOVA with Sidak multiple comparisons. *** *P* < 0.001 by student t-test (A, E, F) and two-way ANOVA with Sidak multiple comparisons test (G-J).

**Figure S2.**
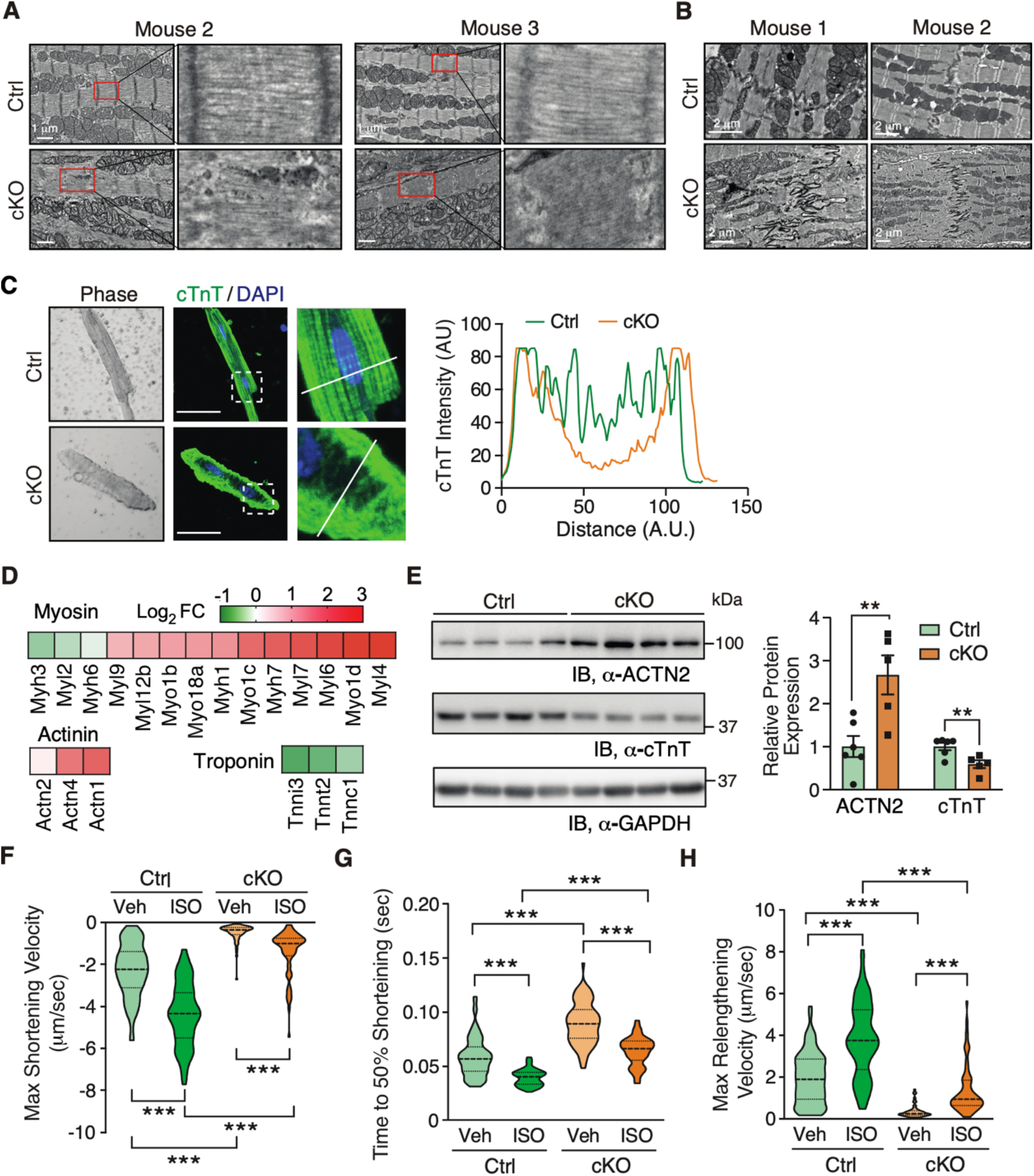
Cardiomyocyte-specific deletion of Fam210a causes myofilament disarray and impairs cardiomyocyte contractility. (**A**) Electron microscopy images of CMs from whole heart tissue sections of the hearts of the other two late-stage *Fam210a* cKO mice (∼65 days post tamoxifen injection). (**B**) Electron microscopy images of two hearts suggest that FAM210A deficiency disrupted intercalated disc integrity between CMs from whole heart tissue sections. (**C**) Immunofluorescence staining of cTnT indicates myofilament disarray in isolated CMs from cKO hearts at ∼65 days post-KO. Right panels: quantification of fluorescence intensity distribution (AU, arbitrary unit) along the white line in the IF image. (**D**) Quantitative mass spectrometry assay shows differentially expressed sarcomeric myosin, actinin, and troponin proteins at the late stage of cKO hearts. (**E**) Western blot confirmed the differential expression of ACTN2 and cTnT in cKO heart lysates. N = 6 for Ctrl and N = 5 for cKO. (**F-H**) FAM210A deficiency attenuated the contractility of isolated CMs with or without ISO stimulation as indicated by Max Shortening Velocity (F), Time to 50% Shortening (G), and Max Relengthening Velocity (H). N = 82/79/77/68 CMs were quantified for Ctrl-Veh, Ctrl-ISO, cKO-Veh, and cKO-ISO from 4 hearts. ** *P* < 0.01; *** *P* < 0.001 by student t-test (E) and Kruskal-Wallis test with Dunn’s multiple comparisons test (F-H).

**Figure S3.**
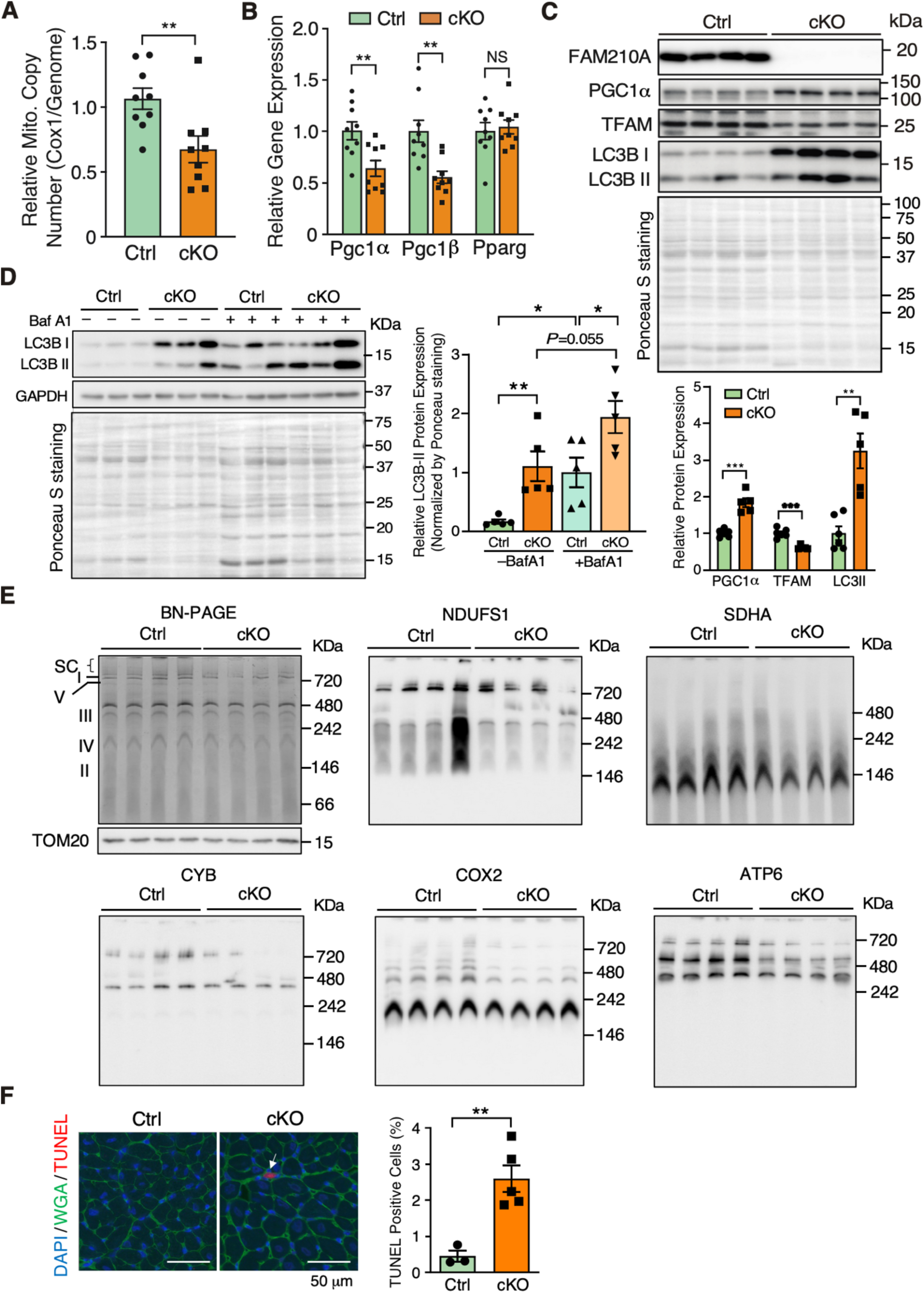
Mitochondrial biogenesis, autophagy, and assembly, and cardiomyocyte apoptosis in CM-specific *Fam210a* knockout hearts at ∼10 weeks post-*Fam210a* KO. (**A**) The mitochondrial copy number was measured by primers targeting the COX1 DNA locus in *Fam210a* cKO hearts (N = 5M + 4F for Ctrl and cKO). The nuclear genomic DNA was used as a normalizer. (**B**) Relative mRNA expression of transcription factors for mitochondrial biogenesis in *Fam210a* cKO and control hearts. β-actin was used as a normalizer. N = 5M + 4F for Ctrl and cKO. (**C**) Western blot measurement of expression of mitochondrial biogenesis and autophagy proteins. The quantification results are shown in the lower panel. N = 6 for Ctrl and N = 5 for cKO. (**D**) Autophagic flux was measured with or without bafilomycin A1 (Baf A1) treatment by western blot for LC3B using the whole heart lysate at the late stage. The quantifications of autophagic flux are shown in the lower panel. (**E**) Blue native PAGE (BN-PAGE) was performed using isolated mitochondria from the late-stage hearts to evaluate the respiratory complex assembly in Ctrl or *Fam210a* cKO hearts. BN-PAGE, followed by immunoblot for ETC component proteins, was used to indicate the molecular weight of complexes, and the same amounts of mitochondria for TOM20 western blot were used as a loading control. (**F**) TUNEL assays for the late-stage Ctrl and *Fam210a* cKO hearts. N = 3 for Ctrl and N = 5 for cKO. Arrow indicates TUNEL positive cell. NS, not significant; * *P* < 0,05; ** *P* < 0.01 by student t-test (A-C, F) and Two-way ANOVA with Turkey’s multiple comparisons (D).

**Figure S4.**
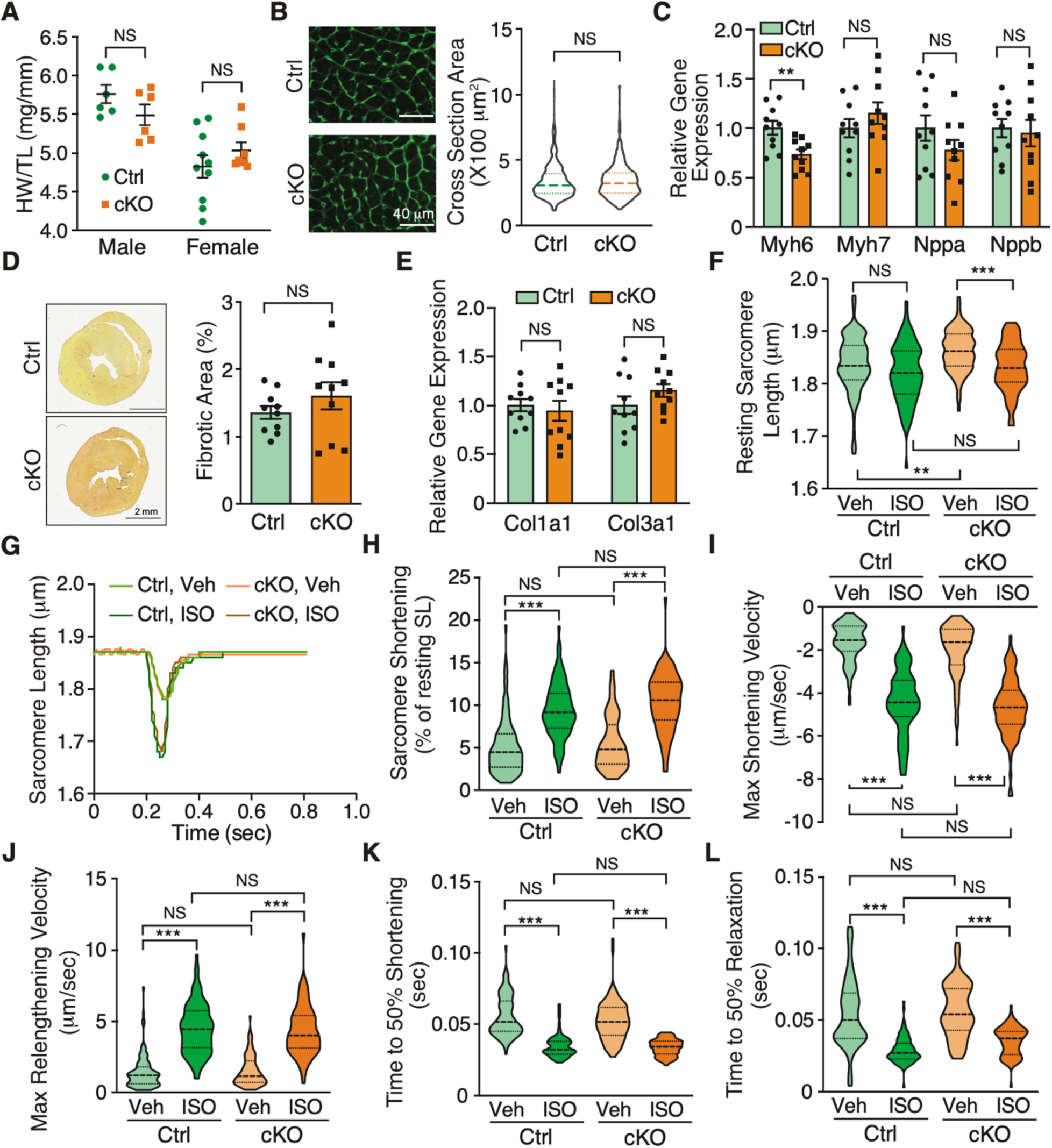
Cardiac function and cardiomyocyte contractility stay unaltered at the early stage of *Fam210a* deficiency. (**A**) The heart weight/tibia length ratio suggests unchanged heart size at 5 weeks post *Fam210a* knockout in male and female mice. N = 6 for male Ctrl and cKO. N = 10 for female Ctrl and N=8 for female cKO. (**B, D**) WGA staining for the cross-sectional area of CMs in B (>750 CMs were quantified from 4 hearts) and picrosirius red staining of collagen deposition in D (N = 5M + 5F for Ctrl and cKO) in the heart of Ctrl and cKO mice at 5-week post Fam210a knockout. (**C, E**) qPCR measurement of hypertrophy and fibrosis marker gene expression at 5-week post-Fam210a knockout. N = 5M + 5F for Ctrl and cKO in C, E. (**F-L**) FAM210A deficiency did not alter the contractility of isolated CMs with or without ISO stimulation at 5-week post-Fam210a knockout as indicated by Resting Sarcomere Length (F), Sarcomere Shortening (G, H), Max Shortening Velocity (I), Max Relengthening Velocity (J), Time to 50% Shortening (K), and Time to 50% Relaxation (L). (**M-O**) FAM210A deficiency did not alter the cytosolic Ca^2+^ in the contraction cycle of isolated CMs with or without ISO stimulation at 5-week post-Fam210a knockout as indicated by Resting Ca^2+^ (M), Fura-2 Ratio (N), and Peak Rate of Ca^2+^ Release (O). N = 100/86/100/86 CMs were quantified for Ctrl-Veh, Ctrl-ISO, cKO-Veh, and cKO-ISO from 4 hearts. NS, not significant; * *P*<0.05; ** *P*<0.01; *** *P*<0.001 by student t-test (C-E), Mann-Whitney test (B), Two-way ANOVA with Tukey’s multiple comparisons test (A), and Kruskal-Wallis test with Dunn’s multiple comparisons test (F, H-O).

**Figure S5.**
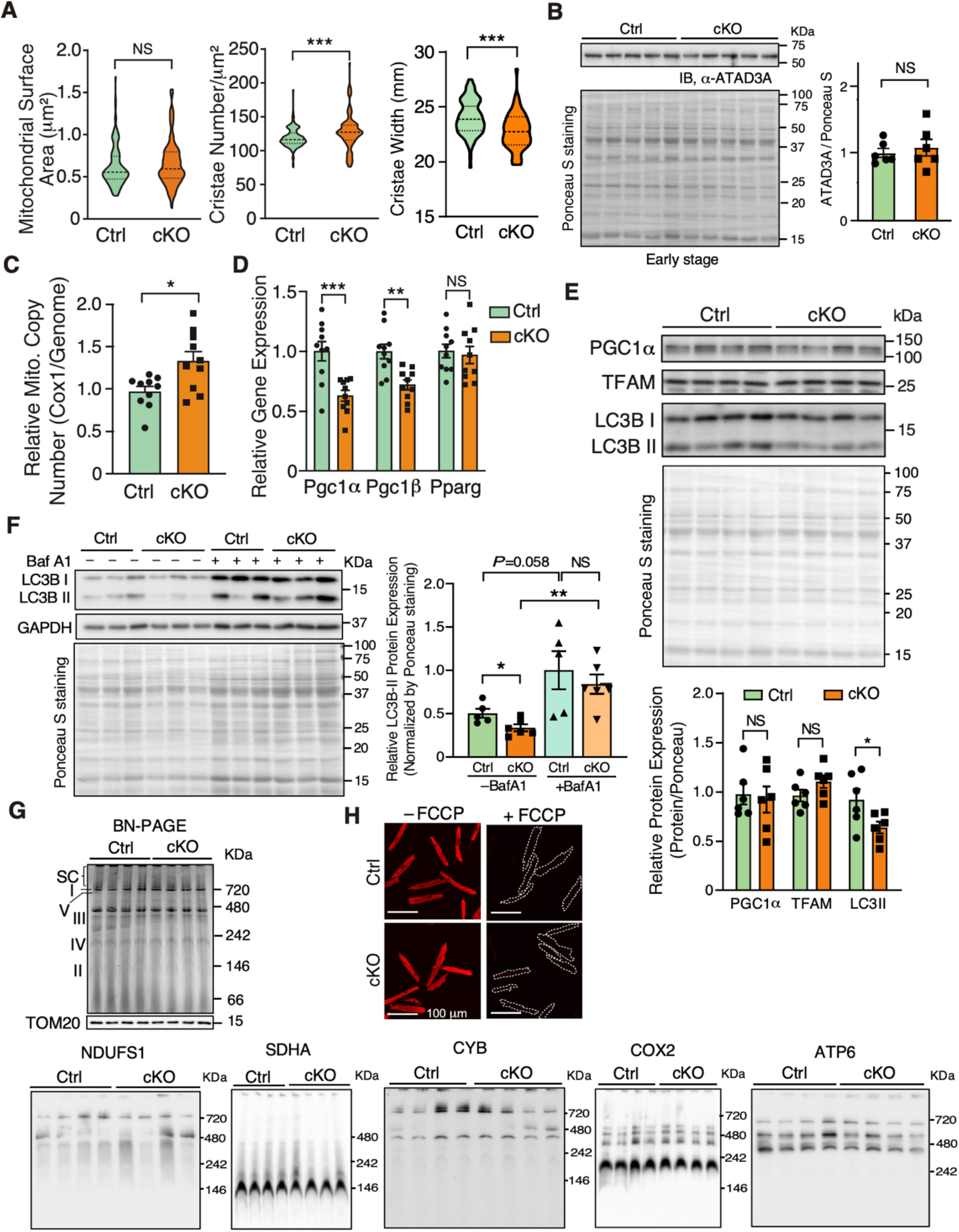
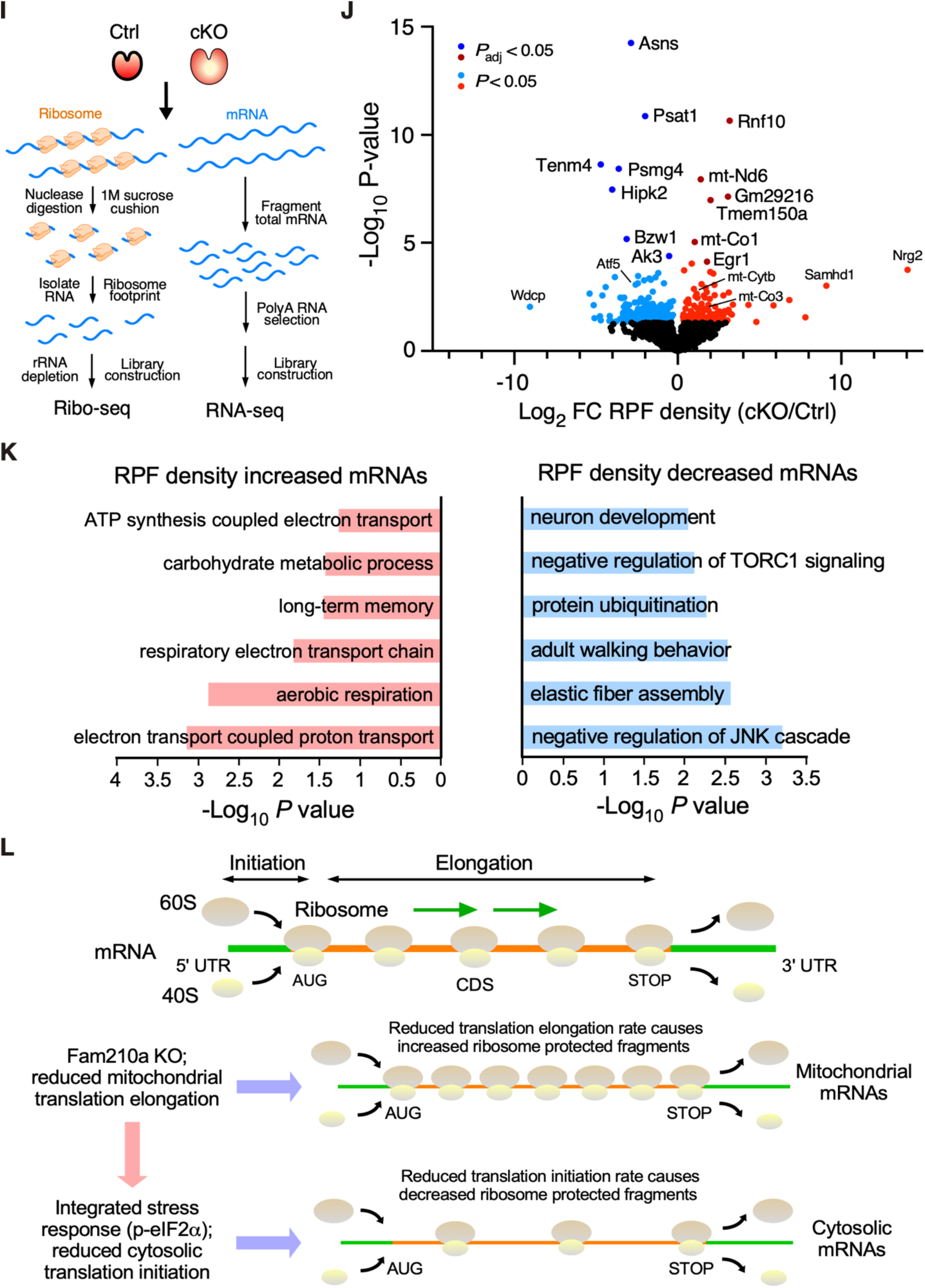
Mitochondrial morphology, copy number, biogenesis, and assembly stay unaltered at 5 weeks post-*Fam210a* knockout in CMs. (**A**) Mitochondrial surface area and cristae number were quantified in electron microscopy images. >100 mitochondria from N = 3 hearts were quantified. Mitochondrial cristae width was quantified from electron microscopy images of the early-stage hearts. N>60 mitochondria were quantified from 3 hearts for each group. (**B**) ATAD3A protein expression was detected by western blot in the early-stage hearts. The quantification is shown in the right panel. N = 6 for Ctrl and cKO at the early stage. NS, not significant. (**C**) Mitochondrial copy number was measured by qPCR using primers targeting the COX1 DNA locus. The nuclear genomic DNA was used as a normalizer. N = 5M + 5F for Ctrl and cKO. (**D**) Relative mRNA expression of transcription factors for mitochondrial biogenesis in *Fam210a* cKO and control hearts at the early stage. β-actin was used as a normalizer (N = 5M + 5F). (**E**) Western blot measurement of mitochondrial biogenesis and autophagy proteins at the early stage. The quantification results were shown in the lower panel (N = 6 for Ctrl and cKO). (**F**) Autophagic flux was measured with or without bafilomycin A1 (Baf A1) treatment by western blot for LC3B using the whole heart lysate at the early stage. The quantifications of autophagic flux were shown in the right panel. (**G**) Blue native PAGE (BN-PAGE) was performed using isolated mitochondria from the early-stage hearts to evaluate the respiratory complex assembly in Ctrl or *Fam210a* cKO hearts. BN-PAGE, followed by immunoblot for ETC component protein, was used to indicate the molecular weight of complexes, and the same amounts of mitochondria for TOM20 western blot were used as a loading control. (**H**) Representative images for Δψ_m_ measurement with the FCCP treatment in Figure 4D. (**I**) Schematic flow chart of ribosome profiling (Ribo-seq). RNA-seq is used as a normalizer to calculate ribosome protected footprints (RPF). (**J**) Ribo-seq analysis of Ctrl and *Fam210a* cKO hearts at the early stage (35 days post-TMX injection). A volcano plot was shown. N = 3 male mice. RPF: ribosome protected fragment. (**K**) Go Ontology analysis of RPF density-upregulated (left panel) and density-downregulated mRNAs (right panel). (**L**) Schematic model of interpretation of the Ribo-seq data for mitochondrial and cytosolic mRNAs. *Fam210a* cKO in CMs may reduce EF-Tu docked on the mitochondrial inner membrane, leading to impaired channeling of aminoacyl-tRNA to the mitoribosome decoding center. This slows down the translocation and movement of mitoribosome (reduced elongation rate) and accumulates more mitoribosomes on mitochondrial-encoded mRNAs, thereby increasing RPF density (more ribosome footprints when normalized by RNA-seq reads). In contrast, *Fam210a* cKO in CMs activates the integrated stress response that inhibits cytosolic translation initiation, leading to reduced loading of cytoribosome on cytoplasmic mRNAs, thereby decreasing RPF density (less ribosome footprints). NS, not significant; * *P* < 0.05; ** *P* < 0.01; *** *P* < 0.001 by student t-test (B-E), Mann-Whitney test (A), and Two-way ANOVA with Turkey’s multiple comparisons (F).

**Figure S6.**
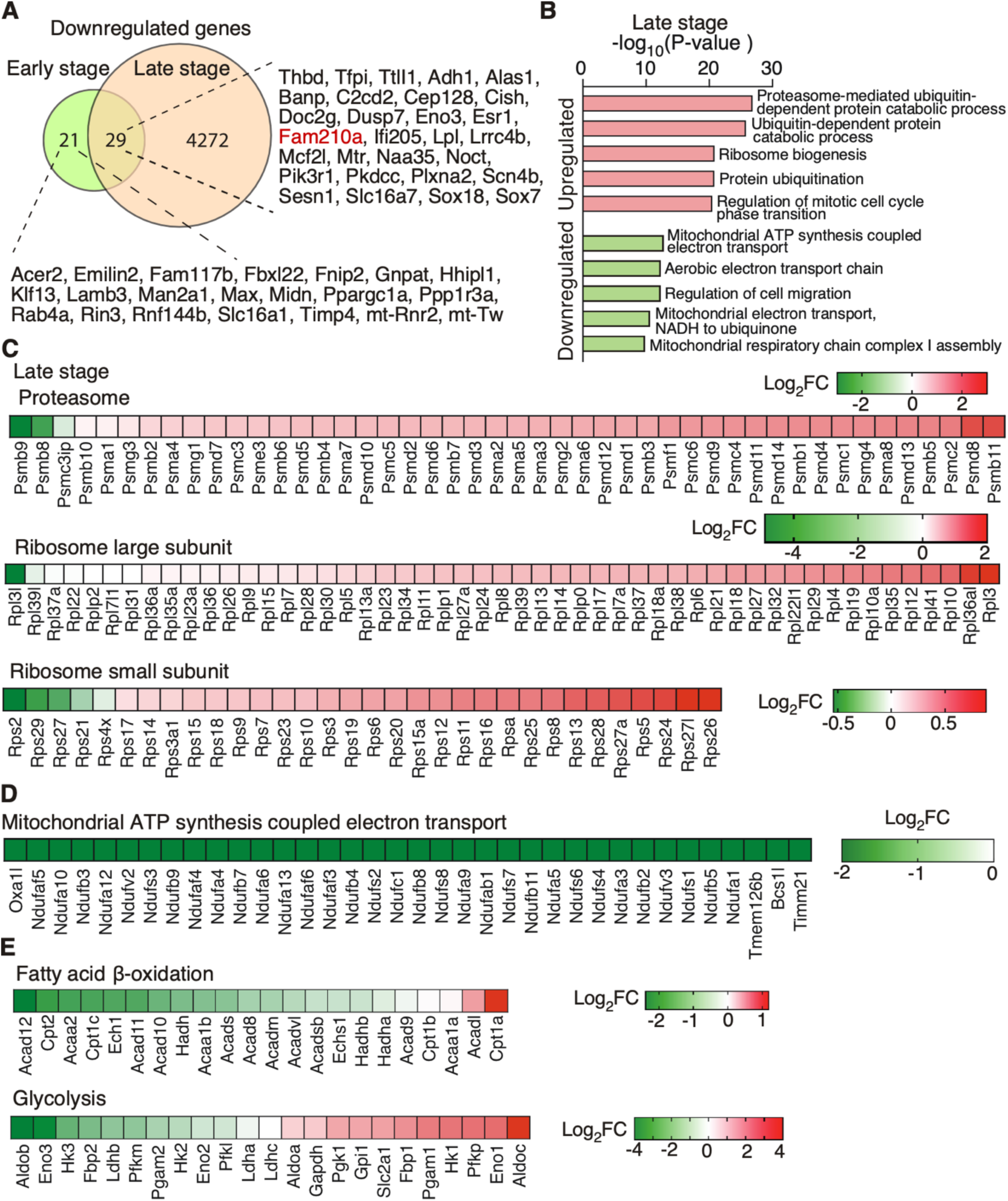
Transcriptomic features at the late stage of FAM210A deficiency. (**A**) The Venn diagram illustrates the overlap of downregulated genes at the transcriptional level in *Fam210a* cKO hearts at the early and late stages. (**B**) GO analyses of biological processes suggest enhanced expression of ribosome and proteasome genes and reduced mitochondrial respiratory chain complex genes at the RNA level. (**C**) The RNA level of most ribosomal and proteasomal genes was induced at the late stage of *Fam210a* cKO hearts. (**D**) mRNA expression of mitochondrial ETC complex components was reduced at the late stage of *Fam210a* cKO hearts. (**E**) Transcriptomic analyses of enzymes in fatty acid β-oxidation and glycolysis suggest a metabolic switch at the late stage of *Fam210a* cKO hearts.

**Figure S7.**
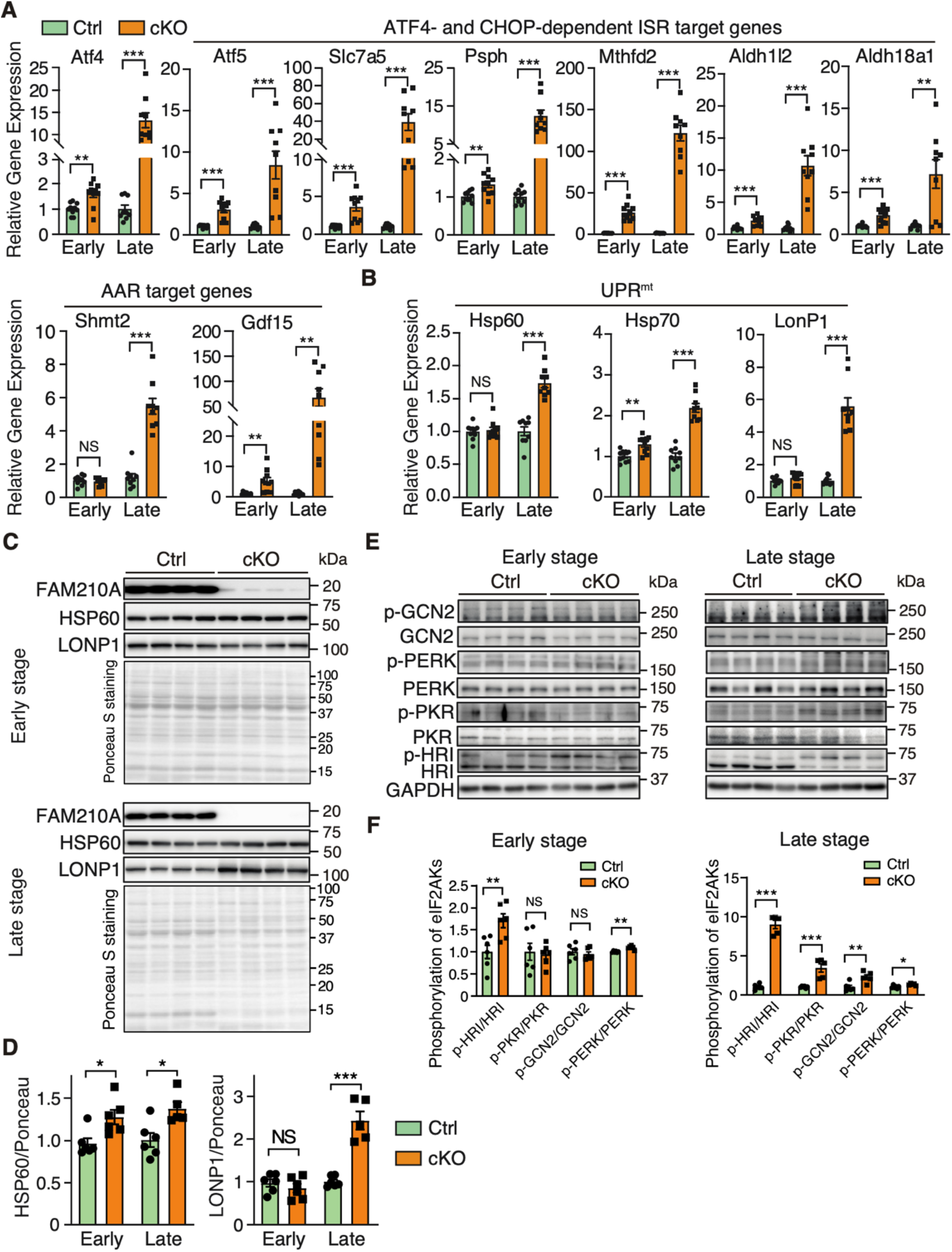
*Fam210a* deficiency persistently activates integrated stress response. (**A**) Relative RNA expression of ATF4 and AAR (amino acid response) target genes shared by ATF4 and CHOP in *Fam210a* cKO hearts at the early and late stages. β-actin was used as a normalizer. N = 5M + 5F for Ctrl and cKO at the early stage. N = 5M + 4F for Ctrl and cKO at the late stage. (**B**) Relative RNA expression of mitochondrial UPR (UPR^mt^) marker genes at early and late stages. N = 5M + 5F for Ctrl and cKO at the early stage. N = 5M + 4F for Ctrl and cKO at the late stage. (**C, D**) Protein expression of UPR^mt^ marker genes at early and late stages (C). Protein expression quantification was shown in D. N = 6 for Ctrl and cKO at the early stage. N = 6 for Ctrl and N = 5 for cKO at the late stage. (**E, F**) Western blot detection of phosphorylation of eIF2α kinases uncover the kinases that activate ISR upon FAM210A deficiency (E). Quantification of phosphorylation of kinases was shown in F. N = 6 for Ctrl and cKO at the early stage. N = 6 for Ctrl and N = 5 for cKO at the late stage. N = 6 for Ctrl and cKO at the early stage. N = 6 for Ctrl and N = 5 for cKO at the late stage. NS, not significant; * *P* < 0.05; ** *P* < 0.01; *** *P* < 0.001 by student t-test (A, B, D, F).

**Figure S8.**
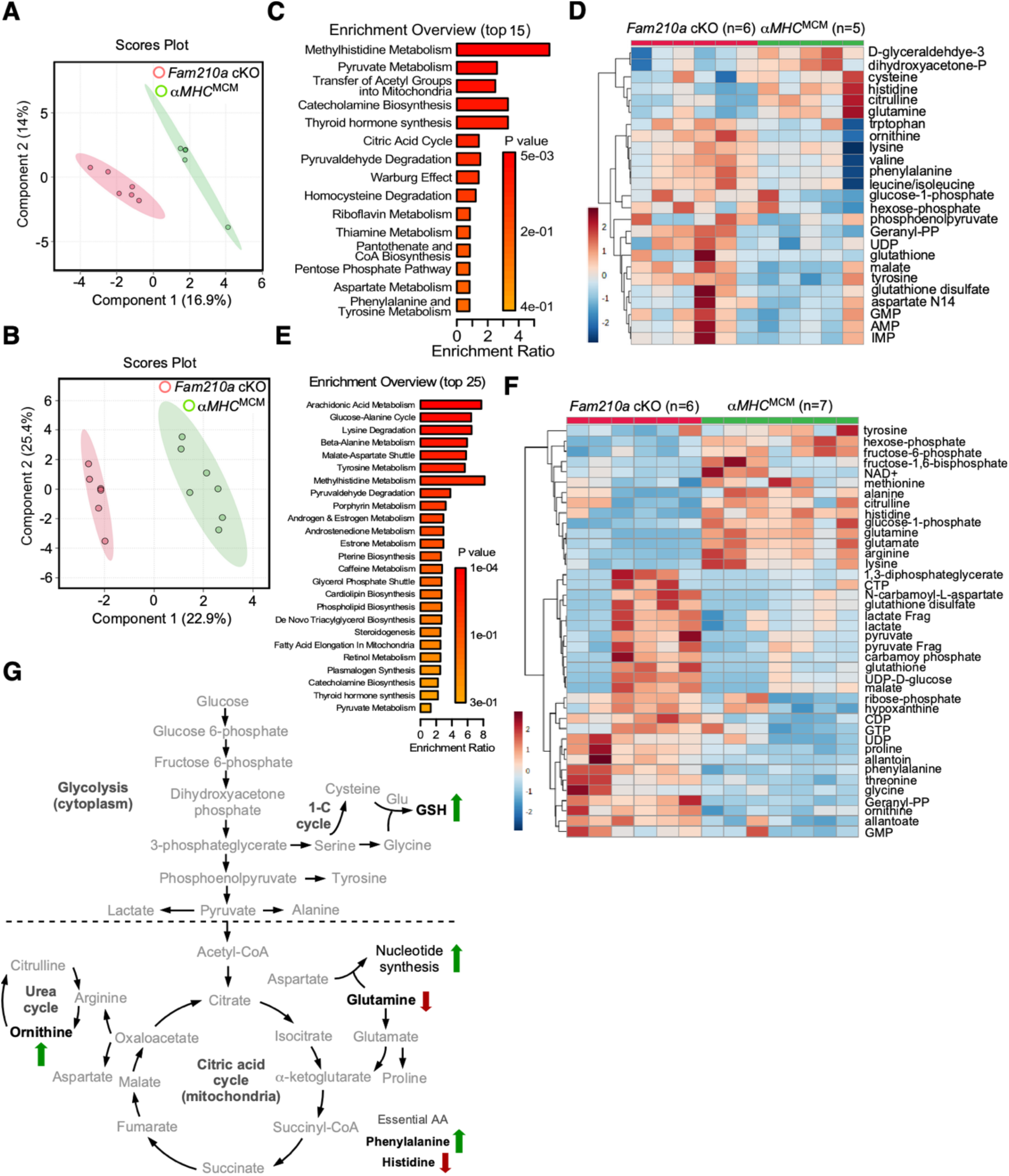
Fam210a loss induces functional alterations in the metabolic profile. (**A**) Principal component analysis of control and *Fam210a* cKO hearts at the early stage (30-day post TMX injection). Note: two dots overlap in control samples. (**B**) Principal component analysis of control and *Fam210a* cKO hearts at the late stage (60-day post-TMX injection). (**C**) Metabolite Set Enrichment Analysis (MESA) of *Fam210a* cKO hearts at the early stage. (**D**) Heatmap of top 25 altered metabolites in *Fam210a* cKO hearts at the early stage. (**E**) Metabolite Set Enrichment Analysis (MESA) of *Fam210a* cKO hearts at the late stage. (**F**) Heatmap of top 40 altered metabolites in *Fam210a* cKO hearts at the late stage. (**G**) Summary of major common metabolites altered in *Fam210a* cKO hearts compared to control hearts at early and late stages. Green arrow: metabolites were increased in quantity in cKO versus control hearts (FC > 1.5 at the early stage and FC > 2 at the late stage); red arrow: metabolites were decreased in cKO versus control hearts (FC < 0.67 at the early stage and FC < 0.5 at the late stage). Metabolites in bold black fonts: *P* < 0.05 by t-tests at the early and late stages.

**Figure S9.**
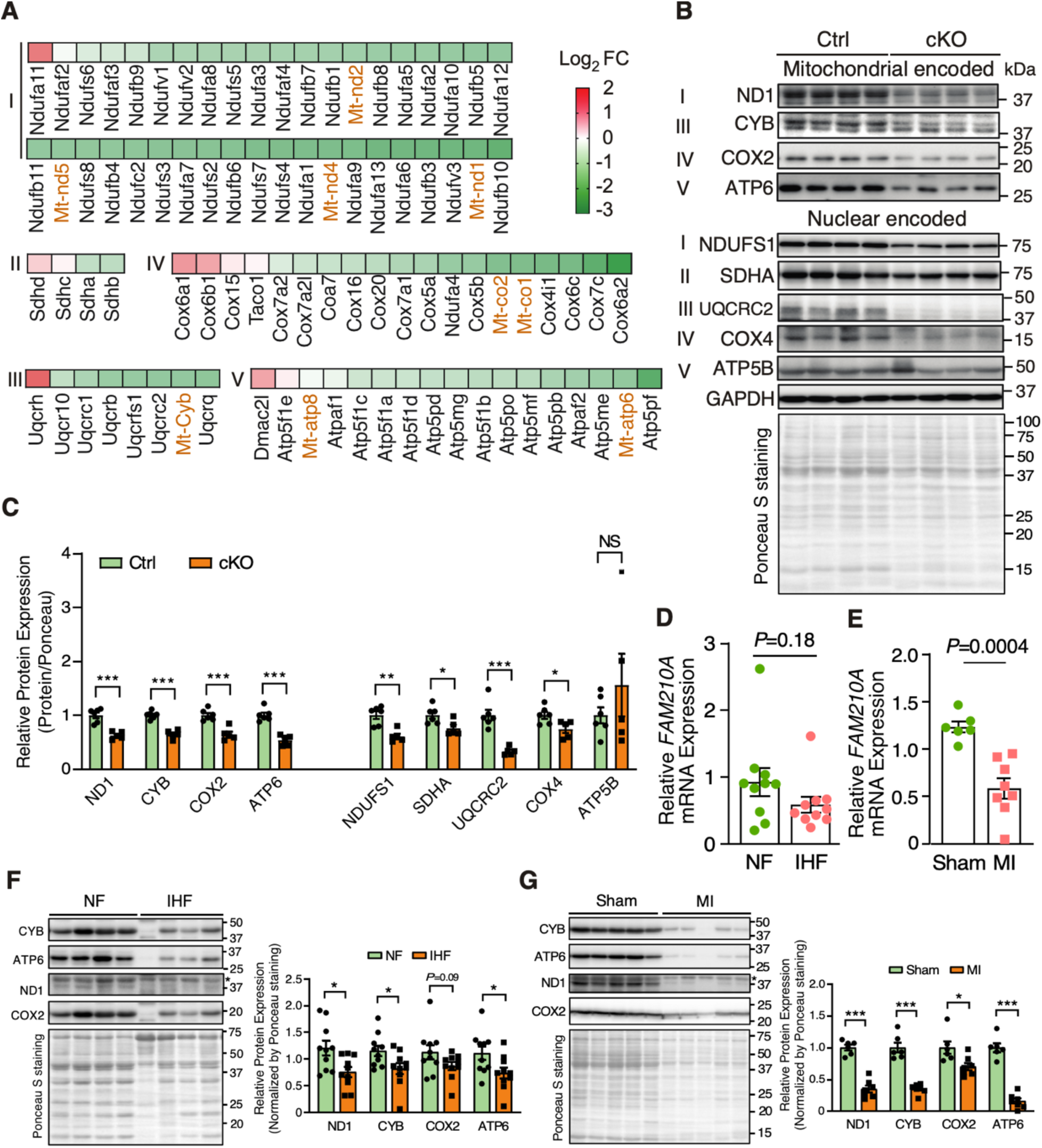
FAM210A depletion causes compromised expression of ETC proteins, and FAM210A is reduced in human and mouse ischemic heart disease. (**A**) Quantitative mass spectrometry analyses of mitochondrial ETC proteins from whole lysates of the late-stage cKO hearts. Red: mitochondrial encoded proteins; Black: nuclear-encoded mitochondrial proteins. (**B**) Western blot detection of the expression of nuclear and mitochondrial encoded ETC proteins in cKO hearts. Ponceau red staining was used as a protein loading control. (**C**) The quantification of the expression of ETC proteins in Figure S9B (normalized by the ponceau red staining). N = 6 for Ctrl and N = 5 for cKO. (**D**) *FAM210A* mRNA expression in human ischemic heart failure (IHF) and non-failing (NF) tissue samples. N = 10 for NF and IHF. 18S rRNA was used as a normalizer. (**E**) *Fam210a* mRNA expression in mouse MI and sham hearts. N = 6 for sham; N = 7 for MI. *Gapdh* mRNA was used as a normalizer. (**F**) Western blot measurement of mitochondrial encoded ETC protein level in human NF and IHF heart samples. *: non-specific band. The quantification results are shown on the right panel. N = 10 for NF and IHF. (**G**) Western blot measurement of mitochondrial-encoded ETC protein levels in mouse Sham and MI hearts. The quantification results are shown on the right panel. N = 6 for Sham and N = 7 for MI. *: non-specific band. * *P* < 0,05; *** *P* < 0.001 by student t-test (C-G).

**Figure S10.**
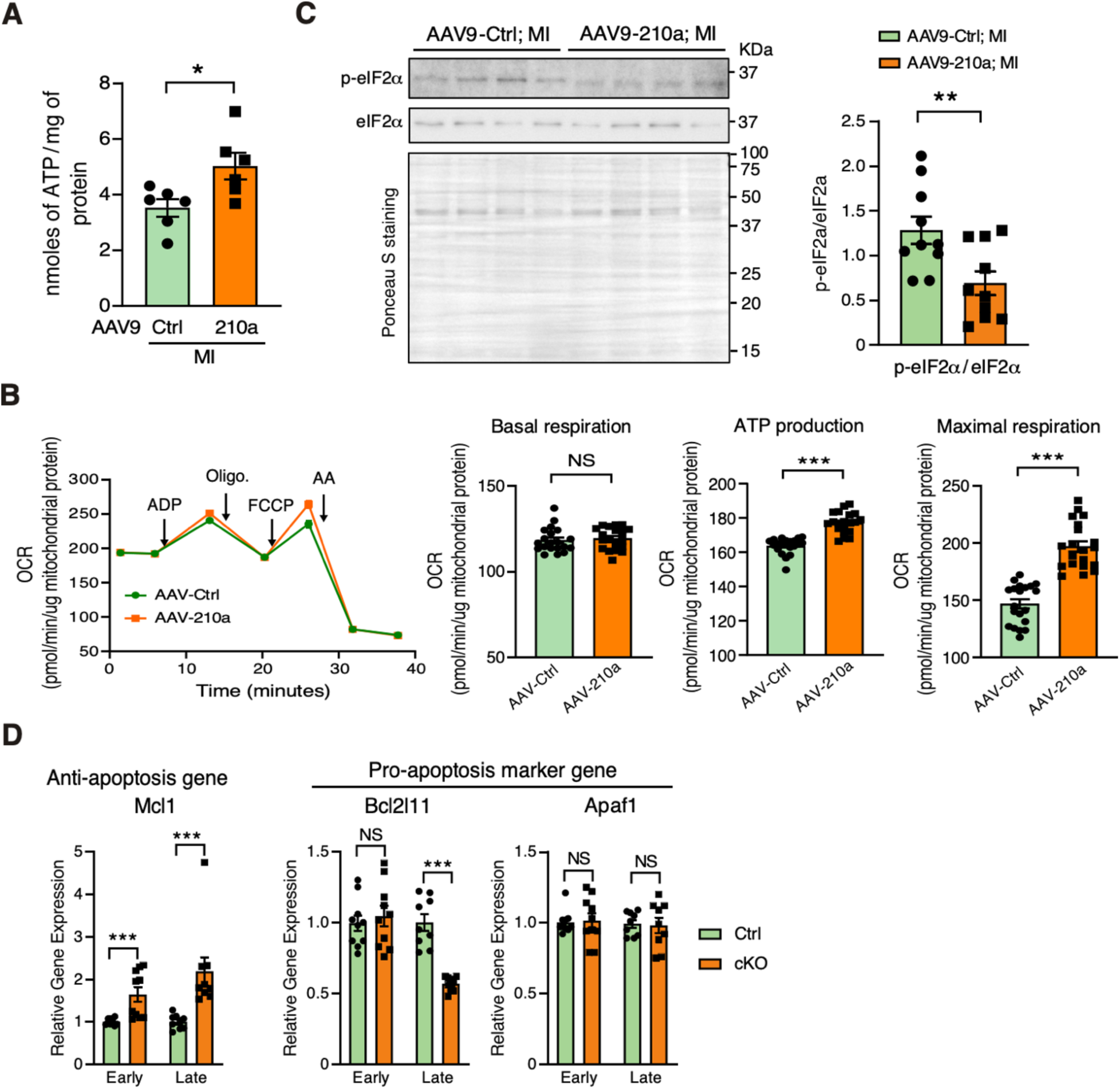
FAM210A overexpression improves mitochondrial function and compromises ISR after myocardial infarction. (**A**) ATP level was determined by ATP determination kit in the whole heart lysate of Ctrl or FAM210A-overexpressing hearts. N = 6 for each group. (**B**) Mitochondrial function was estimated by Seahorse assays using isolated mitochondria from Ctrl or FAM210A-overexpressing hearts 4 weeks after the MI surgery. Oligo.: Oligomycin; FCCP: Carbonyl cyanide-p-trifluoromethoxyphenylhydrazone; AA: Antimycin A. N = 20 biological replicates from 4 CM isolations (2M+2F). (**C**) Western blot detection of phospho-eIF2α and total eIF2α in the whole heart lysate after Fam210a overexpression and 4 weeks MI surgery. Ponceau S staining was used as loading control, and quantifications eIF2α are shown in the right panel. N = 10 for Ctrl (6M+4F) and Fam210a overexpression (3M+7F). (**D**) RNA expression of anti-apoptotic gene *Mcl1* and pro-apoptotic genes *Bcl2l11* and *Apaf1* in the early and late stages of Ctrl and *Fam210a* cKO hearts. NS: not significant; * *P* < 0.05; ** *P* < 0.01; *** *P* < 0.001 by Student t-test (A-D).

## Supplemental tables

**Table S1.** Cardiac function of control and *Fam210a* cKO mice as measured by echocardiography.

**Table S2.** Differentially expressed genes in *Fam210a* cKO hearts compared to control hearts at early and late stages identified by RNA-Seq.

**Table S3**. Differentially expressed proteins in *Fam210a* cKO hearts compared to control hearts at early and late stages identified by mass spectrometry.

**Table S4.** Overlap of significantly dysregulated genes from transcriptomic and proteomic analyses at early and late stages. Gene hits and GO analysis data from Figures 5D, 5F, 5G, and 5H are included in the sub-tables.

**Table S5**. Differentially regulated mRNAs (lncRNAs) at the ribosome-protected fragments (RPF) density level in *Fam210a* cKO hearts compared to control hearts at the early stage identified by ribosome profiling (Ribo-seq) normalized by RNA-seq.

**Table S6.** Overlapped differentially expressed genes from transcriptomic analysis of early and late-stage *Fam210a* cKO hearts. Gene hits and GO analysis data from Figures S6A, S6B, 6B, 6C, and 6E are included in the sub-tables.

**Table S7**. List of genes in existing CHOP ChIP-Seq and ATF4 ChIP-Seq databases and overlapped gene hits with our overlapped differentially expressed genes between early and late stage *Fam210a* cKO hearts from Table S6.

**Table S8.** List of metabolites detected by LC-MS/MS in control and *Fam210a* cKO hearts at the early and late stages. The data were normalized by the median value of each metabolite and analyzed by MetaboAnalyst 5.0.

**Table S9**. Cardiac function was measured by B-mode echocardiography in a myocardial infarction mouse model with FAM210A or control overexpression.

**Table S10.** List of primer sequences for RT-qPCR in this study.

## Materials and methods

### Reagents, antibodies, and cell lines

Tamoxifen (T5648-5g), Isoprenaline (ISO, I5627), Antimycin A (A8674), Oligomycin (O4876), FCCP (C2920), Rotenone (R8875), L-carnitine (C0283), CGP-37157 (C8874) and Digitonin (D141) were purchased from Sigma-Aldrich. BSA-Palmitate Saturated Fatty Acid Complex (5 mM, 29558) and Fura-FF potassium salt (20415) were purchased from Cayman Chemical Company. Dipotassium succinate trihydrate (7156AF) was purchased from AK Scientific. TMRE (T669), MitoSOX Red (M36008), Rhod-2, AM (R1245MP), Thapsigargin (T7459), Insulin-Transferrin Selenium (41400045) were purchased from ThermoFisher Scientific. Ru360 (557440) was purchased from EMD Millipore. Collagenase, Type 2 (LS004177) was purchased from Worthington Biochemical Corporation.

Primary antibodies used in this study include: mouse anti-ACTN2 (A7811; RRID: AB_476766) and rabbit anti-FAM210A (HPA014324; RRID: AB_2668972) were purchased from Sigma-Aldrich. Moue anti-TNNT2 (MA5-12960; RRID: AB_11000742) and rabbit anti-CYB (PA5-43533; RRID: AB_2609847), anti-ATP6 (PA5-37129; RRID: AB_2553922), and anti-ATAD3A (Cat#PA5-66727, RRID: AB_2664945) were purchased from ThermoFisher Scientific. Mouse anti-GAPDH (60004-1-Ig; RRID: AB_2107436), anti-SDHA (66588-1-Ig; RRID: AB_2881948), anti-COX4 (66110-1-Ig; RRID: AB_2881509), anti-MYC-tag (66004-1-Ig; RRID: AB_2881489), anti-FLAG-tag (66008-2-Ig; RRID: AB_2881492), and rabbit anti-UQCRC2 (14742-1-AP; RRID: AB_2241442), anti-ND1 (19703-1-AP; RRID: AB_10637853), anti-CHOP (15204-1-AP; RRID: AB_2292610), anti-MYC-tag (16286-1-AP; RRID: AB_11182162), anti-FLAG-tag (20543-1-AP; RRID: AB_11232216), anti P62/SQSTM1 (18420-1-AP; RRID: AB_10694431), anti-TFAM (22586-1-AP; RRID: AB_11182588), anti-LONP1 (15440-1-AP; RRID: AB_2137152) were purchased from Proteintech Group. Mouse anti-NDUFS1 (sc-271510; RRID: AB_10655669), anti-ATP5B (sc-74549; RRID: AB_1121152) and anti-COX2 (sc-514489; RRID is not available) were purchased from Santa Cruz. Mouse anti-eIF2α (2103S; RRID: AB_836874) and rabbit anti-phopho-eIF2α S51 (3597S; RRID: AB_390740), anti-ATF4 (11815S; RRID: AB_2616025) were purchased from Cell Signaling Technology. Rabbit anti-CYB (A17966; RRID: AB_2861768), anti-ATP6 (A17960; RRID: AB_2861763), anti-MTCO2 (A11154; RRID: AB_2758433), and anti-MT-ND1 (A17967; RRID: AB_2861769) were purchased from Abclonal.

### Human specimens

All human samples of frozen cardiac tissues (n=20), including 10 samples from explanted failing hearts of ischemic heart failure patients and 10 samples from non-failing donor hearts, were acquired from the Cleveland Clinic. The Cleveland Clinic Institutional Review Board approved the study protocol and all participants provided written informed consent. We have been blinded from any clinical data. The human heart tissues were homogenized by Minilys Personal Homogenizer (Bertin Technologies) in TRIzol (ThermoFisher Scientific) or RIPA buffer (ThermoFisher Scientific) for RNA extraction and protein samples, respectively. The mRNA and protein expression of FAM210A was detected by qRT-PCR and immunoblot following the method described below and GAPDH (for immunoblot) and 18S rRNA (for qRT-PCR) were used as the loading control. The experiments conformed to the principles outlined in the Declaration of Helsinki.

### Mice

All mouse experiments were conducted in accordance with protocols approved by University Committee on Animal Resources (UCAR) of the University of Rochester Medical Center (URMC). The mice were housed in a 12:12 hours light: dark cycle in a temperature – controlled room in the animal housing room of URMC, with free access to water and food. The age and gender were indicated below in each section of experiments using mice. The *Fam210a* floxed mice (tm1c, *Fam210a*^flox/flox^) were generated by the Canadian Mouse Mutant Repository (CMMR, http://www.cmmr.ca/) and purchased through the International Mouse Phenotyping Consortium (https://www.mousephenotype.org/). The tamoxifen inducible transgenic mouse line *αMHC*^MerCreMer^ (*αMHC*^MCM/+^) was a gift from Dr. Eric Small lab at URMC. The tamoxifen induced cardiomyocyte specific knockout of *Fam210a* was achieved by crossing *Fam210a*^flox/flox^ *mice with αMHC*^MCM/+^ mice. Age and gender matched *αMHC*^MCM/+^ mice were used as control for *Fam210a* knockout mice. All the mice are on C57BL/6J background.

Anesthetic and analgesic agents used in the study are listed below:

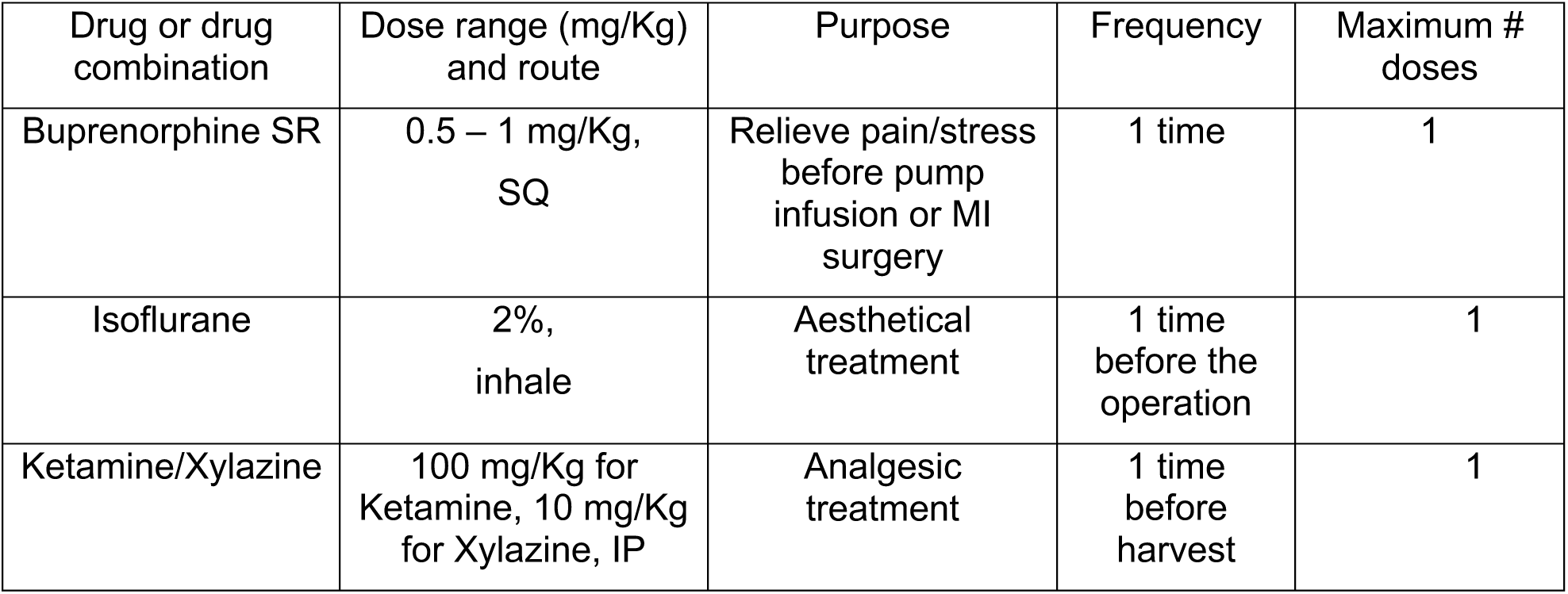

### Tamoxifen administration

Tamoxifen (Sigma) was dissolved in 100% ethanol at concentration of 100 mg/ml in 37°C and corn oil (Sigma) was added to get a final tamoxifen concentration of 10 mg/ml. The control (*αMHC*^MCM/+^) and *Fam210a* cKO (*Fam210a*^flox/flox^, *αMHC*^MCM/+^) mice of both genders at the age of 1-2 months were used for tamoxifen administration. The tamoxifen induction was achived by intraperitoneal injection for four consecutive days at a dosage of 100 μg/g body weight per day for each mouse. The body weight was monitored every week after tamoxifen administration and the knockout efficiency was confirmed by qRT-PCR and Western blot at the end-point of each experiment.

### AAV9-mediated FAM210A overexpression in WT mice

Cardiac Troponin T promoter-driven FAM210A-3xFlag overexpression vector and adeno associated virus serotype 9 (AAV9) were purchased from Vectorbuilder. AAV9-EGFP was used as the control virus (AAV-Ctrl) in our experiments. Control AAV-Ctrl or FAM210A overexpression AAV was administered at postnatal P5-P10 by subcutaneous injection at a dose of 2x10^10^ vg/g following a previous study^1^. Around 2 months after AAV administration, the mice were subjected to LAD ligation to induce myocardial infarction for another 4 weeks, and the heart functions were monitored by echocardiography bi-weekly. At the end-point of 4 weeks post-MI surgery, the hearts were harvested, fixed in 10% neutral buffered formalin, and subjected to paraffin sections with 6 levels. Picrosirius staining and TUNEL assay were performed to detect the scar area and cell death, respectively.

### Terminal deoxynucleotidyl transferase dUTP nick end labeling (TUNEL) assay

The paraffin-embedded sections were deparaffinized by the following steps: xylene (100%) for 2x 5 mins; ethanol (100%) for 2x 5 mins; ethanol (95%) for 1x 5 mins; ddH_2_O for 2x 5 mins. The tissue sections were washed with PBS twice, permeabilized using 0.5% Triton X-100 for 5 mins, and then incubated in the TUNEL reaction mixture (In Situ Cell Death Detection Kit; Sigma, 11684795910) for 1 hr at 37°C in the dark. Finally, the tissue sections were washed with PBS for 3x 5 mins, air dried, and mounted with DAPI-containing antifade medium (Vector Laboratories, H-1500). Images were captured using a confocal microscope (Olympus) and TUNEL positive cells were quantified using the Image J software.

### Echocardiography

Echocardiographic image collection was performed using a Vevo2100 echocardiography machine (VisualSonics, Toronto, Canada) and a linear-array 40 MHz transducer (MS-550D) by the surgical core of Aab Cardiovascular Research Institute (Aab CVRI) at URMC in a blinded manner. Image capture was performed in mice under general isoflurane anesthesia with a heart rate maintained at 500-550 beats/min. Left ventricular systolic and diastolic measurements were captured in M-mode from the parasternal short axis. Echocardiographic images from baseline, 3, 7, 9, 10, and 11 weeks after tamoxifen-induced gene deletion were collected and analyzed from both genders. Hearts were harvested at multiple endpoints in different experiments.

### Measurement of area at risk (AAR) and infarct size (IS)

Left ventricle parasternal long-axis cineloops were acquired via FujiFilm VisualSonics 3100. Wall motion abnormalities were evaluated, identifying IS as completely akinetic dysfunctional areas, AAR as impaired kinetic or semifunctional areas, and healthy tissue as fully kinetic functional areas. Endocardium measurements were made in peak diastole, and infarct size normalized to AAR was quantified following an established method ^2^.

### Wheat germ agglutinin (WGA) staining

The mouse hearts were fixed with 10% neutralized formalin solution and processed for paraffin embedded sections in the Histological Core of Aab CVRI in a blinded manner. WGA staining was used to quantify the size of CMs in the murine heart after tamoxifen-induced gene deletion. The paraffin embedded sections were deparaffinized by the following steps: xylene (100%) for 2x 5 min; ethanol (100%) for 2x 5 min; ethanol (95%) for 1x 5 min; ddH_2_O for 2x 5 min. Antigen retrieval was performed by boiling the deparaffinized section in 10 mM citrate buffer, pH 6.0 and auto fluorescence was quenched by incubation in 3% H_2_O_2_/PBS for 30 min at room temperature (RT). At the staining step, the section dots were marked by Dako pen and stained with 10 μg/ml WGA Alex Fluor-488 (ThermoFisher) for 1.5 hr at RT. Then the slides were washed with PBS for 3x 5 min and finally covered by coverslips with antifade solution (containing DAPI) for imaging. The images were taken in the fluorescence microscope and the Image J software was used to quantify the cell size of CMs.

### Picrosirius red staining

Picrosirius staining was performed to measure the cardiac fibrosis in heart failure models using picrosirius red solution (Abcam) following the manufacturer’s instruction. Briefly, paraffin embedded tissue sections were deparaffinized and incubated in picrosirius red solution at RT for 1 hr. Then, the slides were subjected to 2 washes of 1% acetic acid and 100% ethyl alcohol, and mounted in a mounting medium. Images were captured using the PrimeHisto XE Histology Slide Scanner (Carolina), and the cardiac fibrotic area was quantified from the whole heart images of picrosirius red staining using the Image J software.

### mRNA expression

For heart tissues (human and mouse) or cell samples, the RNA extraction was performed using TRIzol reagent (ThermoFisher) following instructions in the manual and used to quantify the expression of specific genes. Briefly, the tissues were homogenized in TRIzol using Minilys Personal Homogenizer (Bertin Technologies) and placed on ice for 15 min to lyse the tissue.

For the mRNA detection, 1 μg of total RNA was used as a template for reverse transcription using the iScript cDNA Synthesis Kit (Bio-Rad). cDNA was diluted 5-fold and used for detecting the indicated gene expression in each experiment. 18S rRNA or *Gapdh* mRNA was used as a normalization control for mRNA expression. The primer sequences are listed in Table S10. The primers showing only one peak in the melting curve were used in this study. qPCR procedure: 1) initial denaturation at 95°C for 60 sec. 2) 40 cycles of denaturation at 95°C for 10 sec and annealing/extension at 60°C for 45 sec. 3) melt curve analysis by 0.5°C increments at 5 sec/step between 65-95°C.

### Immunoblotting

The heart homogenates were prepared in RIPA buffer (ThermoFisher Scientific) containing a protease inhibitor cocktail (Roche) and phosphatase inhibitor cocktail (Pierce). The tissue lysates were centrifuged at 13,000 rpm at 4°C for 10 min, and the supernatant was saved for the immunoblotting analysis. The protein concentration was measured by Bio-Rad Protein Assay dye, and the lysates were incubated with SDS protein loading buffer (5x stock, National Diagnostics) at 95°C for 10 min. Finally, 20-30 μg of total protein was separated by SDS-PAGE, transferred to PVDF membranes by wet electro-transfer, blocked by 4% milk, and probed with indicated primary antibodies at 4°C overnight. HRP-conjugated secondary antibody and ECL chemical luminescence reagents were used to detect immunoblot signal by the Gel-doc system (Bio-Rad).

### Transmission electron microscopy

Murine hearts at the end-point of control and cKO male mice were flushed with saline containing heparin for 10 seconds. Heart slices were cut and immediately fixed in a flat bottom tube with fixative solution (2.5% glutaraldehyde buffered with 0.1 M sodium cacodylate) that was balanced to RT before use. The samples were placed on a rocking shaker for 1 hr at RT and sent to the Electron Microscopy Share Resource Laboratory for further processing in a blinded manner. Samples were post-fixed in 1% osmium tetroxide, dehydrated in ethanol, transitioned into propylene oxide, and then transferred into Epon/Araldite resin. Tissues were embedded into molds containing fresh resin and polymerized for 2 days at 65°C. 70 nm sections were placed onto carbon-coated Formvar slot grids and stained with aqueous uranyl acetate and lead citrate. The grids were examined using a Gatan 11 mega-pixel Erlansheng digital camera and Digital micrograph software. The morphology of cardiomyocytes and mitochondria were visualized and analyzed in the whole heart tissue sections for phenotyping of *Fam210a* cKO and control mice.

### Adult cardiomyocyte isolation

Adult cardiomyocytes (CMs) were isolated from 2-4 months old male and female mice using a Langendorff perfusion system as previously described^3^. Mice were fully anesthetized via intraperitoneal injection of ketamine/xylazine. Once losing pedal reflex, the mouse was secured in a supine position. The heart was excised and fastened onto the CM perfusion apparatus and perfusion was initiated in the Langendorff mode. Our Langendorff perfusion and digestion consisted of three steps at 37°C: 4 min with perfusion buffer (0.6 mM KH_2_PO_4_, 0.6 mM Na_2_HPO_4_, 10 mM HEPES, 14.7 mM KCl, 1.2 mM MgSO_4_, 120.3 mM NaCl, 4.6 mM NaHCO_3_, 30 mM taurine, 5.5 mM glucose, and 10 mM 2,3-butanedione monoxime), then switched to digestion buffer (300 U/ml collagenase II [Worthington] in perfusion buffer) for 3 min, and finally perfused with digestion buffer supplemented with 40 μM CaCl_2_ for 8 min. After perfusion, the ventricle was placed in sterile 35 mm dish with 2.5 ml digestion buffer and shredded into several pieces with forceps. 5 ml stopping buffer (10% FBS, 12.5 μM CaCl_2_ in perfusion buffer) was added and pipetted several times until tissues disperse readily, and solution turned cloudy. The cell solution was passed through 100 μm strainer. CMs were settled by incubating the cell suspension at 37°C for 30 min. The CMs were resuspended in 10 ml stopping buffer and subjected to several steps of calcium ramping: 100 μM CaCl_2_, 2 min; 500 μM CaCl_2_, 4 min; 1.4 mM CaCl_2_, 7 min. Then the CMs were seeded onto a glass bottom dish (Nest Biotechnology) pre-coated with laminin (ThermoFisher Scientific). Plates were centrifuged for 5 min at 1,000 g at 4°C to increase the adherence, cultured at 37°C for ∼1 hr, and then switched to adult CM culture medium (MEM [Corning] with 0.2% BSA, 10 mM HEPES, 4 mM NaHCO_3_, 10 mM creatine monohydrate, 1% penicillin/streptomycin, 0.5% insulin-selenium-transferrin, and 10 μM blebbistatin for cell culture and downstream assays.

### Immunofluorescence

For the immunofluorescence, the primary cardiomyocyte cells isolated from control and Fam210a cKO mice were fixed using 4% PFA/PBS and washed with PBS for 3x 5 min. The cells were permeabilized by ice-cold 0.5% triton X-100/PBS for 5 min and washed with PBS for 3x 5 min. After blocking with 4% BSA in PBS for 30 min, the cells were incubated with indicated primary antibodies (mouse anti-ACTN2 1:300; mouse anti-Cardiac Troponin T 1:300) in blocking solution (4% BSA in PBS) overnight at 4°C and washed with PBS for 3x 5 min. Then, the cells were stained with the Alex Fluor-488 conjugated secondary antibodies (ThermoFisher Scientific, 1:250) in blocking solution for 45 min and washed with PBS for 3x 5 min. Finally, the cells were co-stained with DAPI for 5 min and kept within a PBS buffer inside before imaging. The images were taken using an Olympus FV1000 confocal microscope and the intensity was measured by the NIH Image J software.

### Measurement of autophagic flux

To evaluate autophagic flux in the heart, bafilomycin A1 (BafA1) was injected at 1.5 mg/kg by intraperitoneal injection (IP) 3 hours before harvesting the heart tissues. The mice were euthanized by ketamine/xylazine mixture, and the hearts were exercised from the mice and lysed using RIPA buffer. Western blot was performed to detect LC3B to indicate the autophagic flux.

### Blue-native PAGE

Blue-native PAGE (BN-PAGE) was performed following the published protocol^4^ with some modifications. The mitochondria were isolated following the instruction of the Mitochondrial Isolation Kit for Tissue (Thermo Fisher Scientific, 89801). The isolated mitochondria were resuspended in a mitochondrial resuspension buffer (225 mM d-Mannitol, 75 mM sucrose, 1 mM EGTA, pH to 7.4), and the mitochondria concentration was measured by Bradford protein assay. Then, 200 ug mitochondria were pelleted in an Eppendorf tube by 10000 g centrifugation for 5 minutes and re-suspended in 1X native loading buffer (Thermo Fisher Scientific, BN2003). Digitonin was added to achieve a digitonin/mitochondria ratio at 6:1, and mitochondria were lysed on ice for 20 minutes. Coomassie blue G-250 dye was added to the samples at a digitonin/dye ratio of 8:1 and glycerol at a final concentration of 1%. Finally, 10 ul of each sample was loaded into each well of 4%-20% native PAGE gel, and electrophoresis was run at 300 v for around 3.5 hours. For the immunoblot after BN-PAGE, the protein complex was transferred to a PVDF membrane using 1X NuPAGE Transfer Buffer (Thermo Fisher Scientific, NP00061) at 40V for 2 hours, and immunoblot was performed using indicated antibodies as described in the methods of western blot part.

### Seahorse assays

Seahorse assay using isolated primary cardiomyocytes from mouse hearts: Mitochondrial respiration rate was measured by the Seahorse assay as previously described^5^ with some modifications. Briefly, adult cardiomyocyte cells were isolated from both genders of early or late stage *Fam210a* cKO and control mice using the Langendorff system described above. The cell viability and yield were determined by Trypan blue and a hemocytometer. Isolated CMs were seeded at 2000/well on Seahorse XF96 V3-PS plates (Agilent Technologies) and incubated in a 37°C incubator with 5% CO_2_ for 1 hr in the adult CM culture medium. Then, the adult CM culture medium was gently replaced with unbuffered DMEM (pH 7.4) supplemented with the following carbon resources: 5 mM glucose, 4 mM glutamine, 0.1 mM pyruvate, 0.1 mM palmitate-BSA, and 0.2 mM L-carnitine and cells were equilibrated at 37°C incubator without CO_2_ for 0.5 hr before measurement. Oxygen consumption rates (OCRs) were measured using an XF96 Extracellular Flux analyzer (Agilent Technologies).

Seahorse assay using isolated mitochondria from mouse hearts: The Seahorse assay using isolated mitochondria was performed following published literature with modifications^6^. The mitochondria were isolated from AAV-Ctrl and AAV-*Fam210a* overexpression hearts after 4 weeks of myocardial infarction following the instruction of the Mitochondrial Isolation Kit for Tissue (Thermo Fisher Scientific, 89801). The isolated mitochondria were resuspended in MAS1 buffer (220 mM d-Mannitol, 70 mM sucrose, KH_2_PO_4_, 5 mM MgCl_2_, 2 mM HEPES, 1mM EGTA, and 0.2% fatty acid-free BSA, adjust pH to 7.2 with KOH) with 10 mM glutamate and 5 mM malate as substrates for complex I-driven respiratory and mitochondria concentration was determined as total protein amount in mitochondria by Bradford protein assay. 2 ug mitochondria were loaded in each well of XF96 cell culture microplate, and oxygen consumption rates (OCRs) were measured using an XF96 Extracellular Flux Analyzer (Agilent Technologies).

### Measurement of mitochondrial superoxide

Mitochondrial superoxide production was measured using the mitochondrial superoxide indicator MitoSOX Red (ThermoFisher Scientific). Briefly, isolated cardiomyocyte cells were plated on glass bottom dishes pre-coated with laminin and equilibrated in a 37°C incubator with 5% CO_2_ for 2 hr in the adult CM culture medium. The cells were loaded with 5 μM MitoSOX Red for 30 min with or without 2 μM antimycin A, washed with pre-warmed PBS, and equilibrated for another 30 min before imaging. The images were obtained using an Olympus FV1000 confocal microscope, and the intensity was measured by the NIH Image J software.

### Measurement of mitochondrial membrane potential

The mitochondrial membrane potential was measured using Tetramethyl rhodamine ethyl ester (TMRE, ThermoFisher Scientific) according to the manufacturer’s protocol. Briefly, isolated cardiomyocyte cells were plated on glass bottom dishes pre-coated with laminin and equilibrated at 37°C incubator with 5% CO2 for 2 hr in the adult CM culture medium followed by 50 nM TMRE loading for 30 min. After incubation, cells were washed with pre-warmed PBS and loaded with the fresh adult CM culture medium. For the normalization, 10 μM FCCP was added into the medium and incubated for 5 minutes before imaging. The images were captured using an Olympus FV1000 confocal microscope, and the intensity was measured by the NIH Image J software. To calculate the mitochondrial membrane potential, the TMRE intensity without FCCP was normalized by the TMRE intensity after FCCP treatment.

### ATP determination assay

The ATP content in the heart tissue was determined using a Bioluminescence kit (Thermo Fisher Scientific A22066) following the manufacturer’s instruction. Briefly, the heart tissue was lysed in RIPA lysis buffer. The protein inside the lysate was precipitated using 1% (W/V) of trichloroacetic acid (TCA) and centrifuged at 10,000 rpm for 5 min at 4 °C. The supernatant was collected and neutralized with 0.1 M Tris-HCl, pH 9.0. Then the equal volume of the supernatant was mixed with the bioluminescent reagent from the kit, and the samples were loaded onto a microplate reader (HTX microplate reader, BioTek instruments). ATP concentration was determined in all the samples and calculated from an ATP standard curve measured in the same microplate. The total protein amount was used to normalize the ATP content in each heart tissue.

### Measurement of cardiomyocyte contractility

The mechanical properties of cardiomyocytes were assessed using a SoftEdge MyoCam system (IonOptix Corporation, Milton, MA)^7^. Cardiomyocytes were placed in a chamber and stimulated with a suprathreshold voltage at a frequency of 0.5 Hz. IonOptix SoftEdge software was used to capture changes in sarcomere length during shortening and relengthening. Cell shortening and relengthening were assessed using the following indices: peak shortening (PS), the amplitude myocytes shortened on electrical stimulation, which is indicative of peak ventricular contractility; time-to-50% shortening and relaxation, the duration of myocytes to reach 50% shortening and relaxation, an indicative of systolic and diastolic duration; and maximal velocities of shortening and relaxation. Intracellular Ca^2+^ was measured using a dual-excitation, single-emission photomultiplier system (IonOptix). Cardiomyocytes were loaded with Fura 2-AM (2 μM) and were exposed to light emitted by a LED lamp through either a 340 or 380-nm filter while being stimulated to contract at a frequency of 0.5 Hz. Fluorescence emissions were then detected.

### Analysis of mitochondrial translational activity by polysome profiling

Mitochondria were isolated from control and *Fam210a* cKO hearts following the instruction of the Mitochondria Isolation Kit for Tissue (Thermo Fisher Scientific, 89801). The mitochondrial polysome profiling was performed as previously described^8^. Briefly, mitochondria isolated from control and *Fam210a* cKO hearts were lysed on ice for 20 min using mitochondrial lysis buffer (260 mM sucrose, 100 mM KCl, 20 mM MgCl_2_, 10 mM Tris-Cl [pH 7.5], 1% Triton X-100, 2 mM DTT, protease inhibitor cocktail without EDTA [Roche] and 4 U/mL RNase inhibitor [NEB]). Lysates were cleared by centrifugation at 12000 g for 10 min at 4°C. An equal amount of A260 absorbance from each sample was loaded to 10%-30% sucrose gradient solution and centrifuged at 29,000 rpm for 4 hours in a Sorvall Surespin 630 (17 ml) rotor. After centrifugation, 8 fractions were collected from each sample by Density Gradient Fractionation System (BRANDEL). Total RNA was extracted from the same fraction volume with Trizol LS (ThermoFisher Scientific). Renilla luciferase mRNA from *in vitro* transcription was used as RNA spike-in loading control for qRT-PCR measuring the distribution of mitochondria-encoded mRNAs across each fraction.

### RNA-Seq NGS data processing and alignment

Total RNA extracted from a mouse heart left ventricle tissues was treated with DNase I (NEB) to remove potential genomic DNA in the RNA samples. The DNase I treated RNA samples were purified with phenol:chloroform: isoamyl alcohol and then subjected to RNA-Seq at the Genomic Research Center of URMC. Raw reads generated from the Illumina HiSeq2500 sequencer were demultiplexed using bcl2fastq version 2.19.0. Quality filtering and adapter removal are performed using Trimmomatic version 0.36^9^ with the following parameters: "TRAILING:13 LEADING:13 ILLUMINACLIP: adapters.fasta:2:30:10 SLIDINGWINDOW:4:20 MINLEN:15". Processed/cleaned reads were then mapped to the *Mus musculus* reference genome (GRCm38, mg38) with STAR_2.5.2b^10^ given the following parameters: "—twopassMode Basic --runMode alignReads --genomeDir $^11^ --readFilesIn ${SAMPLE} --outSAMtype BAM SortedByCoordinate-outSAMstrandField intronMotif --outFilterIntronMotifs RemoveNoncanonical". The subread-1.5.0^12^ package (featureCounts) was used to derive gene counts given the following parameters: “-s 2 -t exon -g gene_name” and the gencode M12 gene annotations. Differential expression analysis and data normalization were performed using DESeq2-1.16.1^13^ with an adjusted p-value threshold of 0.05 within an R-3.4.1^14^ environment. A batch factor was given to the differential expression model in order to control for batch differences. Gene ontology and KEGG pathway enrichment analyses were performed using the Enrichr webtool^15^.

### Metabolomics analysis of the whole heart

The liquid chromatography-tandem mass spectrometry (LC-MS/MS) based metabolomics were performed as previously described^16^. The hearts from *Fam210a* cKO and control mice were perfused with saline and snap-frozen in liquid nitrogen. The frozen hearts were grounded in a cooled pestle and mortar into power. The heart powder was serially extracted in 80% aqueous methanol. The extracts were evaporated to dryness under nitrogen gas and resuspended in 50% aqueous methanol. Metabolites were separated by reverse phase HPLC (Shimadzu) and identified by a triple-quadrupole mass spectrometer (Thermo Quantum Ultra) running in a negative mode with selected-reaction monitoring (SRM)-specific scans. The LC-MS/MS data were analyzed using a publicly available mzRock machine learning tool kit (http://code.google.com/p/mzrock/), which automates SRM/HPLC feature detection, grouping, signal-to-noise classification, and comparison to known metabolite retention times. Data for each run were median-normalized, and analyzed and visualized using the MetaboAnalyst tool^17^.

### Ribosome profiling

Ribosome-protected footprint (RPF) RNA preparation: Ribosome profiling was performed following the published protocol^18^ with some modifications. For heart homogenization, the mice were anesthetized with the ketamine/xylazine mixture, and the hearts were quickly excised out, cut out atrium, washed with cold PBS containing 100 μg/ml cycloheximide (CHX), and snap frozen in liquid nitrogen. The ventricular tissues were homogenized at 4 °C using a Minilys Personal Homogenizer (Bertin Technologies) in 900 μl ribosome lysis buffer (150 mM NaCl, 5 mM MgCl_2_, 1 mM DTT, 20 mM Tris-HCl, pH 7.4 with 100 μg/ml CHX and 25 U/ml DNase I). For complete lysis, the samples were kept on ice for 10 minutes and passed through a 26½ gauge needle 5 times. Then the samples were centrifuged at 15,000 g for 5 minutes, and the supernatant was immediately used for further steps. 100 μl of the supernatant was lysed in 1 ml Trizol for total RNA extraction, and the remaining supernatant was digested using RNase I (N6901K, LGC, Biosearch Technologies) at a concentration of 0.5 U/μg RNA at room temperature with gentle agitation for 1 hour. After digestion, the monosomal fraction was pelleted by ultracentrifuge on a 1M sucrose cushion using Sorvall S120-AT2 rotor at around 530,000 g for 1 hour. The ribosome-protected RNA was extracted from the pellet using 500 μl Trizol following the manufacturer’s instructions, and ribosome footprints with a size range of 17-34nt were further purified by electrophoresis on 15% TBE-urea gel. Finally, the ribosome footprints RNA was sent to the Genomic Research Center of URMC and subjected to small RNA sequencing.

Small RNA Library Preparation for RPF: RNA concentration and quality were determined using the NanopDrop 1000 spectrophotometer and Agilent Bioanalyzer 2100 methods, respectively. rRNA depletion was first performed utilizing the NEBNext rRNA Depletion kit (New England Biolabs), following manufacturer’s specifications. To optimize retention of small RNA fragments (ribosome protected footprints), the final sample purification step was modified with the addition of isopropanol to the bead binding step. The rRNA-depleted RNA was then immediately utilized as input for sequencing library construction with the NEBNext Small RNA Library Prep kit (New England Biolabs), following the manufacturer’s protocol. Importantly, undiluted 3’ and 5’ adaptors were utilized for ligation steps, as specified by manufacturer due to the small size of the RNA. Following 3’ adaptor ligation, RT primer incubation, and 5’ adaptor ligation, cDNA was synthesized, and final library amplification was performed with 12 cycles of PCR. Libraries were isolated from the PCR reaction mix using the Qiagen MinElute PCR Reaction Cleanup kit, quantity and quality were assessed using a Qubit fluorometer (ThermoFisher) and Bioanalyzer (Agilent), respectively, and sequenced on an Illumina NextSeq 550 High output flowcell generating single-end reads of 75 nt.

Bioinformatic analysis of RPF density (demultiplexing, quality control, alignment, and analysis): For Ribo-seq analysis, raw reads generated from the Illumina basecalls were demultiplexed using bcl2fastq version 2.19.1. Quality filtering and adapter removal are performed using FastP version 0.23.1 with the following parameters: "--length_required 7 --cut_front_window_size 1 --cut_front_mean_quality 3 --cut_front --cut_tail_window_size 1 --cut_tail_mean_quality 3 --cut_tail-y –r".1 rRNA reads were filtered out, and the Processed/cleaned reads were then mapped to the GRCm39/gencode M31 or GRCh38/genecode38 (Mouse OR Human) reference using STAR_2.7.9a with the following parameters: "—twopass Mode Basic --runMode alignReads --outSAMtype BAM Unsorted – outSAMstrandField intronMotif --outFilterIntronMotifs RemoveNoncanonical –outReadsUnmapped Fastx --windowAnchorMultimapNmax 2-- -- seedSearchStartLmax 15 -n 3 -l 5".2,3 Genelevel read quantification was derived using the subread-2.0.1 package (featureCounts) with a GTF annotation file (GRCm39/gencode M31 or GRCh38/gencode42) and the following parameters for stranded RNA libraries "-s 2 -t exon -g gene_name" -M --fraction". Fractional counts were rounded to whole numbers for translation efficiency calculations. RiboDiff was used to calculate translation efficiency from the RiboSeq data and the Novogene mRNA expression data.

RNA-seq analysis (normalizer for Ribo-seq) was performed by Novogene company following the procedure (library construction, quality control, and sequencing): mRNA was purified from total RNA using poly-T oligo-attached magnetic beads. After fragmentation, the first strand cDNA was synthesized using random hexamer primers, followed by the second strand cDNA synthesis. The library was ready after end repair, A-tailing, adapter ligation, size selection, amplification, and purification. The library was checked with Qubit and real-time PCR for quantification and a bioanalyzer for size distribution detection. Quantified libraries were pooled and sequenced on Illumina platforms, according to effective library concentration and data amount.

### Statistical Analysis

All quantitative data were presented as mean ± SEM, and analyzed using Prism 8.3.0 software (GraphPad). The Kolmogorov-Smirnov test was used to calculate the normal distribution of the data. For normally distributed data, an unpaired two-tailed Student t-test was performed for the comparisons between two groups and one-way or two-way ANOVA with Tukey’s multiple comparisons test for the comparisons between more than three groups. For not normally distributed data, non-parametric Mann-Whitney test was performed for the comparisons between two groups, and Kruskal-Wallis test with Dunn’s multiple comparisons test for the comparisons between more than three groups. The Log-rank test was performed for the comparisons of survival rate, and χ^2^ test was used for comparisons of cell number counting data. Two-sided p values <0.05 were considered to indicate statistical significance. Specific statistical methods were described in the figure legends.

## Notes

### Competing Interest Statement

The authors have declared no competing interest.

